# Genomic signatures of climate-driven (mal)adaptation in an iconic conifer, the English yew (*Taxus baccata* L.)

**DOI:** 10.1101/2025.05.29.656833

**Authors:** Thomas Francisco, Maria Mayol, Elia Vajana, Miquel Riba, Marjana Westergren, Stephen Cavers, Sara Pinosio, Francesca Bagnoli, Maurizio Marchi, Filippos A. Aravanopoulos, Anna-Maria Farsakoglou, Ivan Scotti, Bruno Fady, Giovanni G. Vendramin, Juliette Archambeau, Andrea Piotti, Santiago C. González-Martínez

**Author notes:** Thomas Francisco, Maria Mayol, Elia Vajana, Miquel Riba, Marjana Westergren, Stephen Cavers, Sara Pinosio, Francesca Bagnoli, Maurizio Marchi, Filippos Aravanopoulos, Anna-Maria Farsakoglou, Ivan Scotti, Bruno Fady, Giovanni G Vendramin, Juliette Archambeau, Andrea Piotti, Santiago C. González-Martínez.

## Abstract

The risk of climate maladaptation is increasing for numerous species, including trees. Developing robust methods to assess population maladaptation remains a critical challenge. Genomic offset approaches aim to predict climate maladaptation by characterising the genomic changes required for populations to maintain their fitness under changing climates. In this study, we assessed the risk of climate maladaptation in European populations of English yew (*Taxus baccata*), a long-lived tree with a patchy distribution across Europe, the Atlas Mountains, and the Near East, where many populations are small or threatened. We found evidence suggesting local climate adaptation by analysing 8,616 SNPs in 475 trees from 29 European *T. baccata* populations, with climate explaining 18.1% of genetic variance and 100 unlinked climate-associated loci identified via genotype-environment association (GEA). Then, we evaluated the deviation of populations from the overall gene-climate association to assess variability in local adaptation or different adaptation trajectories across populations and found the highest deviations in low latitude populations. Moreover, we predicted genomic offsets and successfully validated these predictions using fitness proxies assessed in plants from 26 populations grown in a comparative experiment. Finally, we integrated information from current local adaptation, genomic offset, historical genetic differentiation and effective migration rates to show that Mediterranean and high-elevation *T. baccata* populations face higher vulnerability to climate change than low-elevation Atlantic and continental populations. Our study demonstrates the practical use of the genomic offset framework in conservation genetics, offers insights for its further development, and highlights the need for a population-centred approach that incorporates additional statistics and data sources to credibly assess climate vulnerability in wild plant populations.

## Introduction

Genetic adaptation is one of the main mechanisms for long-term persistence of organisms in a changing environment (Aitken *et al*. 2008). For genetic adaptation to occur, individuals must exhibit heritable phenotypic differences, and these differences must affect their survival and/or reproduction (i.e., be related to their fitness; Darwin 1859; Mayr 1982). While evidence for local adaptation in plants (Leimu and Fischer 2008) and animals (Hereford 2009) is common, it can only occur, or be maintained, when the selective pressures are greater than other evolutionary forces such as genetic drift or gene flow (Savolainen *et al*. 2013). Random genetic drift can, purely by chance, promote mutations that are not beneficial but rather neutral or even deleterious, potentially undermining local adaptation (Yeaman and Otto 2011). Gene flow can hinder adaptation by introducing maladaptive alleles, or promote it by spreading beneficial alleles across populations and increasing overall genetic diversity (Joron *et al*. 2011; Blanquart *et al*. 2013; Savolainen *et al*. 2013; Deacon and Cavender-Bares 2015).

The predicted intensification of climate change in the near future poses serious threats to all biodiversity levels (Díaz *et al*. 2019). Rapid environmental changes expose species and populations to novel conditions leading to a phenotypic mismatch, which may result in a reduction in fitness (Chen *et al*. 2011; Brady *et al*. 2019). A growing body of evidence shows range contractions and shifts as serious threats to biodiversity conservation, particularly affecting sessile organisms that will most likely not be able to adapt and disperse at the same rate as the predicted climate change (Zhu *et al*. 2012; Dyderski *et al*. 2018; Zu *et al*. 2021). Additionally, fragmentation often leads to smaller population sizes, which increases the likelihood of genetic drift and inbreeding, while limiting gene flow (Young *et al*. 1996; Cheptou *et al*. 2017). Therefore, isolated populations may be more vulnerable to climate change due to reduced adaptive capacity and migration potential (Leimu *et al*. 2010). Thus, the development of reliable methods to study and predict maladaptation is becoming urgent, in particular for long-lived, sessile keystone species with highly-fragmented ranges, such as some trees (e.g., the English yew, *Taxus baccata* L.).

To quantify levels of maladaptation in populations under changing climate, Fitzpatrick and Keller (2015) proposed an approach based on genomic data and climate projections, known as the genomic (or genetic) offset. This increasingly popular approach relies on calculating the magnitude of genetic change required to maintain the current gene-environment relationships under novel climates, assuming that populations are locally adapted (Bernatchez *et al*. 2024). Several methods have been used to compute genomic offsets: e.g., latent factor mixed models (LFMMs; Gain and François 2021), gradient forest (GF; Fitzpatrick and Keller 2015) or redundancy analysis (RDA; Capblancq and Forester 2021), and have been applied to a variety of study systems: crops (Aguirre-Liguori *et al*. 2019; Rhoné *et al*. 2020), trees (Capblancq *et al*. 2020b; Theraroz *et al*. 2024; Archambeau *et al*. 2025; Bonnier *et al*. 2025), invertebrates (Adam *et al*. 2022; Bourret *et al*. 2024), amphibians (Hung *et al*. 2023), birds (Bay *et al*. 2018; Smith *et al*. 2021) and, recently, mammals (Hoste *et al*. 2024; McLennan *et al*. 2025).

Recent studies have shown that genomic offset predictions can be validated using simulation (Láruson *et al*. 2022) or empirical (Bay *et al*. 2018; Fitzpatrick *et al*. 2021; Archambeau *et al*. 2025) approaches. However, to date, most validations have relied on the same data used to train the model. For this reason, some authors have highlighted the need to use independent datasets to train and test genomic offset models in order to avoid model overfitting (e.g., Rellstab *et al*. 2021 ; Lotterhos 2024b). Furthermore, the majority of the evaluation studies using empirical data produced contrasting results depending on the fitness proxy used (Fitzpatrick *et al*. 2021; Lind *et al*. 2024a; Archambeau *et al*. 2025). For example, in trees, Lind *et al*. (2024a) found that height was more associated with genomic offset predictions than mortality for Jack pine (*Pinus banksiana* Lamb.) and Douglas fir (*Pseudotsuga menziesii* Mirb.), while Archambeau *et al*. (2025) found the opposite pattern for maritime pine (*Pinus pinaster* Aiton). These contrasting results suggest that the most appropriate fitness proxies may vary across species, even for those with similar life-history traits. Taken together, these findings highlight the need for more case studies across species and ecological scenarios, as well as evaluation data that use a variety of fitness proxies (e.g., variables related to reproductive, phenological or ecophysiological traits), before adopting the genomic offset approach to support population conservation or management guidelines.

Although the genomic offset framework is promising and offers a variety of methods, several assumptions, common to all methods, need to be carefully considered (see Ahrens *et al*. 2023). These include: (i) all populations are equally locally adapted and (ii) the genotype-environment relationships can be extrapolated across space and will remain valid in the future, thus neglecting the adaptive capacity of populations (Rellstab *et al*. 2021; Ahrens *et al*. 2023). Populations deviating from overall gene-climate relationships may be particularly common in species with fragmented distributions as some populations would not or only partially exchange genes with other populations (Deacon and Cavender-Bares 2015). To date, these key assumptions remain largely untested (but see the RONA framework; Rellstab *et al*. 2016). Moreover, a number of additional processes can exacerbate climate maladaptation (e.g., inbreeding depression) or mitigate it (e.g., beneficial gene flow; Aguirre-Liguori *et al*. 2021; Rellstab *et al*. 2021; Chen *et al*. 2022; but see Lachmuth *et al*. 2024). In particular in trees, the overall effect of gene flow is expected to facilitate the evolutionary change required by populations to adapt to new climatic conditions, as they typically exhibit long-distance gene flow (Kremer *et al*. 2012). Several recent studies have interpreted together (Capblancq *et al*. 2020b; Lazic *et al*. 2024) or combined (Barratt *et al*. 2024) various sources of information with genomic offset to provide more integrative climate maladaptation or vulnerability predictions, but the integration of key processes such as gene flow is still largely missing.

English yew (*Taxus baccata* L.) is a long-lived, dioecious, wind-pollinated and animal-dispersed conifer tree species, with a fragmented distribution across Europe, the Atlas Mountains and the Near East, from sea level up to 2,200 m a.s.l. Many populations of *T. baccata* are small and isolated, and they are threatened by fires, browsing and, in some cases, current or past overexploitation and long-term climate variability (Thomas and Polwart 2003; Mysterud and Østbye 2004; Ruprecht *et al*. 2010; Benham *et al*. 2016; Bach *et al*. in revision). The species is found in a wide range of climatic conditions, including high-elevation, Atlantic, continental, and Mediterranean climates (Thomas and Polwart 2003). Cold temperature and drought appear to be among the main limiting factors explaining, respectively, its northern (Thomas and Polwart 2003) and southern (Gegechkori 2018; but see Linares 2013) distribution. Several studies have described overlaying levels of population genetic structure in *T. baccata* at different geographical scales. Indeed, while Mayol *et al*. (2015) found two to three main gene pools at the rangewide geographical scale, studies at local and regional spatial scales found strong population genetic structure and marked genetic differentiation even between populations in close spatial proximity (e.g., less than 500 metres in Dubreuil *et al*. 2010; see also González-Martínez *et al*. 2010; Chybicki *et al*. 2011). Different lines of evidence suggest local adaptation to climate in this species, including phenotypic differences observed under common garden conditions (Mayol *et al*. 2020), as well as genomic signatures of natural selection in defense-related genes (Burgarella *et al*. 2012) and climate-related genes (Mayol *et al*. 2020). Demographic decline is thought to have begun a long time ago in several parts of the *T. baccata* range (e.g., around 300,000 years ago in Iberian populations; Burgarella *et al*. 2012) and may have intensified more recently (particularly over the past 4,000 years; Thomas and Polwart 2003). Climate change is expected to exacerbate this decline, particularly for Mediterranean populations at the warm and arid range limit, and for those in other sensitive environments, such as high-elevation mountain regions, where climate-induced stress is already being observed and/or is predicted to be particularly strong in the near future (Mendoza *et al*. 2009; Knight 2022).

The main objectives of this study are i) to investigate patterns of local adaptation to climate in *T. baccata*, ii) to evaluate the potential of genomic offset to predict population maladaptation under novel climatic conditions for a species with a highly-fragmented distribution, and iii) to gain insight into the potential vulnerability of *T. baccata* populations to future climate by linking information on the current degree of local adaptation, the historical capacity of gene flow and the predicted future climate maladaptation. To meet these objectives, we first used climatic and genomic data from 29 populations across the species’ European range to identify gene-climate relationships, as well as candidate climate-associated loci; second, we computed the distance between the observed and predicted genomic composition to identify populations deviating from average gene-climate relationships (subsequently called ‘genomic discrepancy index’); and third, we calculated historical effective migration and genetic differentiation between populations to determine potential trends in gene flow capacity in the near future, assuming conservatism of historical patterns. Finally, we calculated genomic offsets using two methods and evaluated their predictions using phenotypic traits related to growth, growth phenology, reproductive phenology and drought/temperature tolerance, measured in plants from 26 populations grown in a comparative experiment.

## Materials and methods

### Sampling and genotyping

Genomic data were obtained for 501 trees from 29 populations covering nearly the entire range of *T. baccata* in Europe (Figure 1 and Table S1). A first dataset involving 25,726 Single Nucleotide Polymorphisms (SNPs) from 120 trees sampled in 12 natural populations was obtained from Mayol *et al*. (2020). A subset of these SNPs (7,405 SNPs) was used as target SNPs to genotype 11 additional populations (202 individuals) using Single Enrichment Primer Technology (SPET) by Farsakoglou *et al*. (in revision), data available at Data INRAE public repository (doi: 10.15454/7COC1A). In addition, 179 trees from six populations in Slovenia and Italy were genotyped using the same SPET technology. For the latter, DNA was extracted using the NucleoSpin Plant II kit (Macherey Nagel). SNP calling followed GATK best practices (version 3, with –filterExpression ‘MQ < 40 || MQrank > 12.5 || DP < 7.0 || Q < 50.0 || QD < 1.5 || FS > 60.0 ||ReadPosRanksum > -8’; Auwera and O’Connor 2020).

**Figure 1.**
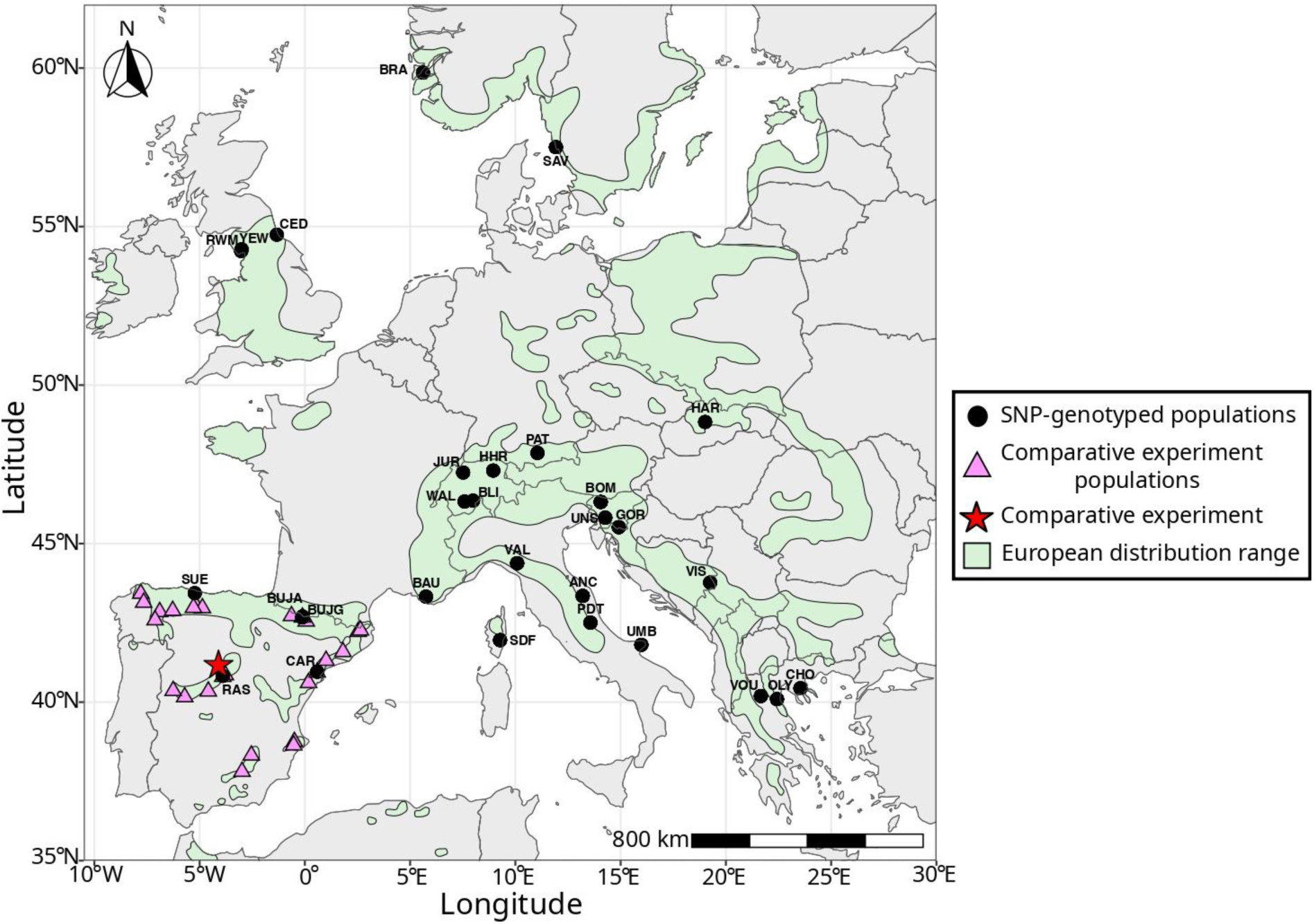
Map showing the two sets of *Taxus baccata* populations used for genomic offset (and related indexes) estimation and validation. In green, the species’ European range (EUFORGEN; Caudullo *et al*. 2017). The black dots represent the 29 SNP-genotyped populations, while the pink triangles indicate the 26 populations from which material was sourced to establish the comparative experiment in Central Spain (clonal bank) used for empirical evaluation of genomic offset models. Population labels correspond to those in Table S1.

Additionally, we filtered for allele balance (between 0.2 and 0.8) using the vcfR R package (Knaus and Grünwald 2017). A subset of three individuals that were genotyped by both Mayol *et al*. (2020) and Farsakoglou *et al*. (in revision) was used to ensure reproducibility across experiments. Low-quality SNPs and those with inconsistent calls across experiments were discarded, resulting in a final genomic dataset of 11,367 SNPs for 490 trees.

From these data, two datasets were produced for specific analyses (see Table S2). For the first dataset, we filtered for missing data > 15% (for both individuals and SNPs) while retaining low-frequency alleles, resulting in 8,252 SNPs for 452 individuals. This set was used for population genetic structure analyses, as they are sensitive to high missing data rates and removal of low-frequency SNPs (Linck and Battey 2019). For the second dataset, used for all other analyses (except for estimating historical gene flow, see below), we filtered for missing data > 30% (for both individuals and SNPs) and removed all SNPs with a minor allele count lower than 20 (i.e., twice the number of individuals in the smallest population before filtering; corresponding to a minor allele frequency of 2.11%), as low-frequency alleles can lead to high false-positive rates, resulting in 8,616 SNPs for 475 individuals (ranging from 8 to 31 individuals per population). Then, we imputed missing data in this dataset by using the most common genotype within each gene pool identified by a STRUCTURE clustering analysis (Pritchard et al. 2000; see below).

Finally, a dataset consisting of 4,993 individuals from 238 populations genotyped with seven nuclear microsatellites (nuSSRs) was retrieved from Mayol *et al*. (2015). This dataset was used to estimate historical gene flow using EEMS (see below and Table S2). Analyses were also performed based only on populations with at least ten trees sampled (4,658 individuals, 176 populations).

### Climatic data

Climatic data at high resolution (30 arc-seconds, approximately one kilometre at the equator) were obtained from the Climate Downscaling tool (ClimateDT, available at www.ibbr.cnr.it/climate-dt; Marchi *et al*. 2024). The 1901-1950 time interval, which likely reflects the long-term climatic conditions to which current *T. baccata* populations are adapted, was defined as the reference period. To predict maladaptation to future climate, we used climatic projections for the 2041-2070 time interval from five global climate models (GCMs; Table S3) under the Shared Socio-economic Pathway SSP3-7.0 (AR6) associated to a severe climate forcing scenario (Fujimori *et al*. 2017).

To identify climatic variables potentially involved in *T. baccata* adaptation, we applied four selection steps. First, we preselected climatic variables based on their relevance according to previous studies and/or the species’ ecology (Melzack and Watts 1982; Moir 1999; Thomas and Polwart 2003; Mayol *et al*. 2015, 2020; Gegechkori 2018; Sánchez-Martínez *et al*. 2021; Cedro 2023). Second, we performed forward selection on the preselected variables to identify those explaining most of the genomic variation among populations. For this, we used the ordiR2step function from the vegan R package (Oksanen *et al*. 2025), ranging from a null model without explanatory variables to a full model with all the explanatory variables. The stopping criteria was the significance of the variable in increasing the model’s *R*², using a *p*-value threshold of 0.05 and 1,000 permutations. Twenty runs were performed to ensure robustness (Table S4). Third, we removed climate variables with Pearson’s correlation greater than |0.75|, keeping, when possible, the most informative climate variables identified during the forward selection. Fourth, we removed additional climatic variables to achieve variance inflation factors (VIF) below 10, following James *et al*. (2021). The final six selected climatic variables were: mean annual temperature, mean diurnal range temperature, temperature seasonality, mean temperature of the driest quarter, mean annual precipitation and precipitation seasonality (Table 1).

**Table 1.** Description of the final set of six climatic predictors used in genotype-environment association (GEA) and genomic offset analyses. “OrdiR2step model” corresponds to climatic predictors identified by the forward selection as explaining most of the genomic variation among populations.

| Variable | Abbreviation | Relevance |
| --- | --- | --- |
| *Bio1*: Mean Annual Temperature (°C) | Annual_Tc | Thomas and Polwart (2003); Gegechkori (2018); Mayol *et al.* (2019); Ahmadi *et al.* (2020) |
| *Bio2*: Mean Diurnal Range (°C) | Diurnal_range_Tc | OrdiR2step model |
| *Bio4*: Temperature Seasonality (standard deviation x100) | Tc_Seasonality | Cedro and Iszkulo (2011); Ahmadi *et al.* (2020); OrdiR2step model |
| *Bio9*: Mean Temperature Driest Quarter (°C) | Tc_driest_quarter | Moir (1999); Cedro and Iszkulo (2011); OrdiR2step model |
| *Bio12*: Mean Annual Precipitation (mm) | Annual_P | Moir (1999); Mayol *et al.* (2019) |
| *Bio15*: Precipitation Seasonality (coefficient of variation) | P_Seasonality | Ahmadi *et al.* (2020) |

To establish how well our SNP data depicted rangewide climatic pressures, we visually compared the climatic envelope of the 29 SNP-genotyped populations with the rangewide one based on all 298 locations with confirmed presence of *T. baccata* used in this study (i.e., including the locations genotyped with nuSSRs in Mayol *et al*. 2015; Figure S1). The climatic envelopes were built by extracting, for each location, the values for the six climate predictors identified above and combining them using PCA (FactoMineR; Lê *et al*. 2008). In addition, we compared the climatic envelope of these 29 populations with that of the 26 populations from the comparative experiment used to validate the genomic offset models.

### Phenotypic traits

Phenotypic data were obtained from plants from 26 natural populations grown in a comparative experiment (a clonal bank established in 1992) under common garden conditions. Only three of the populations (RAS, CAR and BUJA) in the comparative experiment were also in the SNP dataset, and probably sampled different individual trees, therefore we considered both sets of populations to be nearly independent. The comparative experiment was located in Valsaín, in the foothills of the Sierra de Guadarrama mountains in central Spain (40.9106°N, 4.0125°W, 1,140 m a.s.l). Phenotypic traits were measured several times over the years for over 250 trees (see details in Mayol *et al*. 2020). We selected growth, growth phenology, reproductive phenology and drought/temperature tolerance as relevant fitness components for climate adaptation, based on previous studies and the ecology of the species, as explained below (see also Table S5 and Mayol *et al*. 2020).

Growth is commonly considered a proxy for fitness in trees (Fitzpatrick *et al*. 2021; Capblancq *et al*. 2023). Individual growth was approximated using shoot volume measured as the volume of the leading current-year shoot from the two longest stems, averaged between the stems and over four years (2009-2012).

Phenological traits often follow latitudinal/altitudinal clines, suggesting climate-related selective pressures that may lead to adaptation (Alberto *et al*. 2013; Mayol *et al*. 2020; Silvestro *et al*. 2023). This is especially relevant for *T. baccat*a, because some populations occur at high elevation or latitude, where they are exposed to frost events, whilst others occur at low elevation or latitude, and are exposed to drought during the growing season or benefited by milder climate (coastal Atlantic populations). Growth phenology was estimated from the mean ratio of shoot length before summer (May-June) to the total shoot length at the end of the growing season (October-November), over three consecutive years (2010-2012; referred to as shoot elongation). Reproductive phenology was estimated over the same period by estimating the ratio of open male strobili to the total number of male strobili in several branches during the main period of pollen dispersal (see details in Mayol *et al*. 2020).

Finally, drought/temperature tolerance traits have garnered increasing interest for evaluating the potential of populations to adapt to warming climates (Keller *et al*. 2011). Previous studies in *T. baccata* have shown that drought may be one of the limiting factors in the southern part of the species’ range (Thomas and Polwart 2003; Gegechkori 2018). We used the mean leaf thickness, an ecophysiological trait associated with drought and thermal tolerance, measured with a digital calliper (to the nearest 0.01 mm) on the two-five largest leaves of the leading shoot at the end of the growing year, over four years (2009-2011, 2021).

For all traits, we calculated best linear unbiased predictors (BLUPs) to estimate population-level phenotypic values, accounting for covariates influencing trait variation when needed. As the traits showed nearly normal distributions, we used Gaussian mixed-effects models as follows:

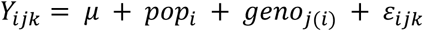

where 𝑌_𝑖𝑗𝑘_is the average trait value across years of measurement for individual *k* of genotype *j* in population *i*, 𝜇 is the overall phenotypic mean, 𝑝𝑜𝑝_𝑖_ and 𝑔𝑒𝑛𝑜_𝑗(𝑖)_ are the population and genotype (nested in population) random intercepts, respectively, and 𝜀_𝑖𝑗𝑘_ are the residuals of the model. For the growth model, stem length was used as a covariate (fixed effect) to account for differences in age between trees. BLUPs used in this study correspond to the mean of the posterior distribution of 𝑝𝑜𝑝_𝑖_.

All mixed-effects models were fitted within a Bayesian framework using Markov chain Monte Carlo (MCMC) methods implemented in the MCMCglmm R package (Hadfield 2010). Because of lack of prior knowledge, we used weakly informative priors (inverse-Wishart with V=1 and ν=0.002) for all random effects and residuals (Hadfield 2010). Each model ran for 1,500,000 iterations, with a 50,000- iteration burn-in to stabilise the chains and a thinning interval of 500 iterations to reduce autocorrelation. Four independent MCMC chains were run for each model and chain convergence was assessed using the Gelman-Rubin criteria (Gelman and Rubin 1992) (Table S5).

### Population genetic structure and historical gene flow

Population genetic structure analyses were conducted using principal component analysis (PCA), as implemented in the vegan R package (Oksanen *et al*. 2025), and the STRUCTURE Bayesian clustering approach (version 2.3.4; Pritchard *et al*. 2000). STRUCTURE models were run for a number of clusters (*K*) ranging from one to ten, with ten independent MCMC runs for each *K*, using 500,000 iterations each, including a burn-in period of 100,000. The most likely number of genetic clusters was estimated from averaged values for each *K,* using the *ΔK* method (Evanno *et al*. 2005).

Historical gene flow patterns were investigated based on the genetic differentiation (*F*_ST_) of each population relative to a common predicted ancestral gene pool, calculated using Bayescan software version 2.1 (Foll and Gaggiotti 2008). Potential areas of historical permeability or barriers to gene flow across the range of *T. baccata* were further identified by estimating effective migration surfaces (EEMS, Petkova *et al*. 2016). This analysis evaluates the consistency of molecular data with expectations under an isolation by distance (IBD) model (see details in Table S7 and in Supplementary Material).

### Patterns of local adaptation to climate

#### Genetic variance partitioning

To disentangle the influence of demographic history, geography and climate on population allele frequencies in *T. baccata*, we used a combination of redundancy analysis (RDA) and partial redundancy analysis (pRDA), following Capblancq and Forester (2021). We first ran a full RDA model with all predictors to estimate the total proportion of genetic variance they explain. Second, we ran three pRDA models to estimate the proportion of genetic variance explained separately by demographic history, geography and climate, while partialling out the effects of the others.

Population demographic history was accounted for by using the first two PCs (explaining 24.5 % of the variance) of the PCA performed on the SNP data not filtered for minor allele count. As proxies for geography, we used all positive correlation axes from a spatial autocorrelation analysis (distance-based Moran’s eigenvector maps, dbMEMs) computed with the adespatial R package (Dray 2016).

Finally, the selected climate variables (Table 1) were used as climate predictors. All the RDA and pRDA models, and the PCA, were performed using the vegan R package (Oksanen *et al*. 2025).

#### Climate-related candidate SNPs

SNPs potentially involved in climate adaptation were identified using six genotype-environment association (GEA) methods: RDA, pRDA, LFMM, BayPass, and Gradient Forest, with or without correction for population structure (see Table 2, Figure 2 and additional Methods in Supplementary Material). These methods have contrasting assumptions and performance under different scenarios, and combining their results helps reduce false positive and negative rates.

**Figure 2.**
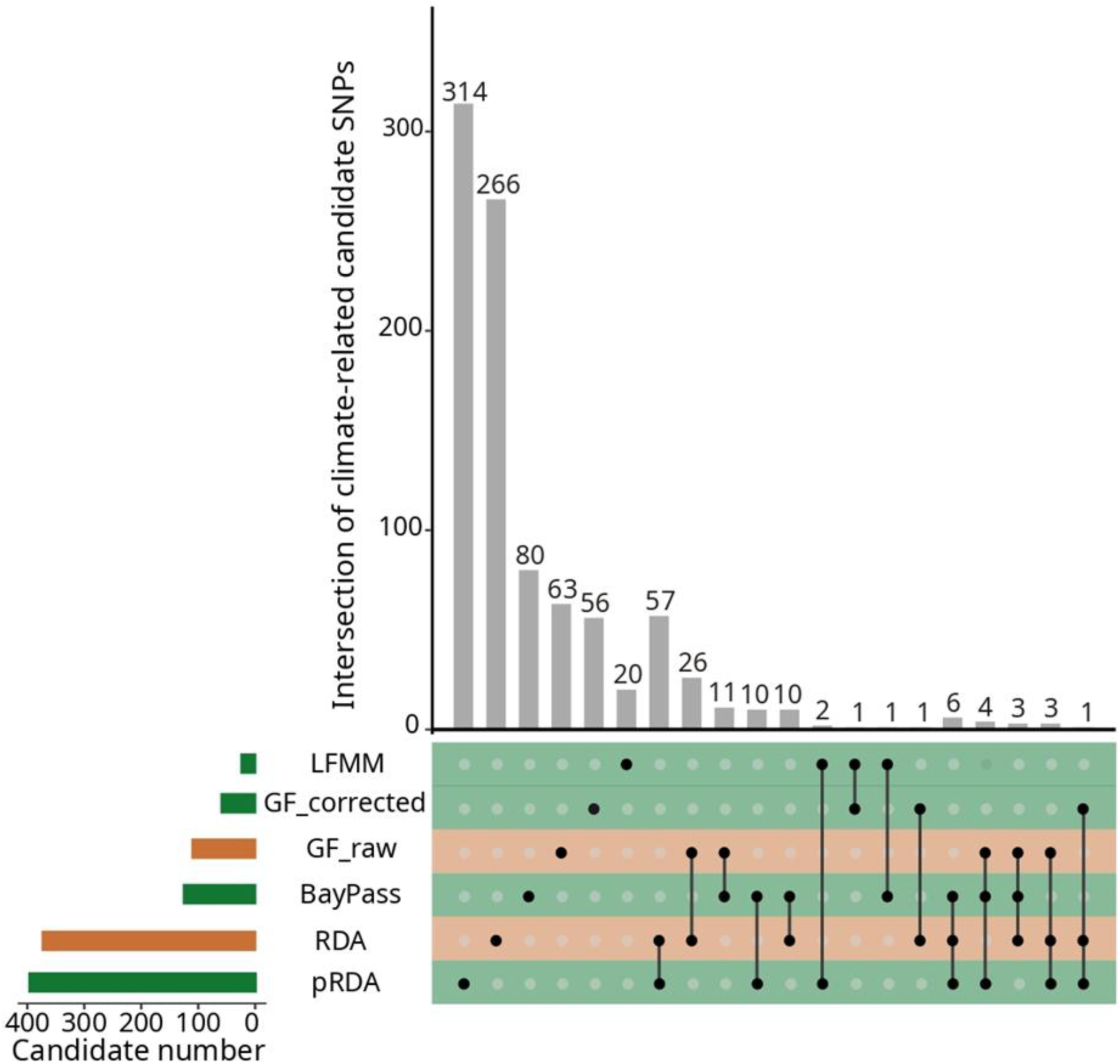
UpSet plot for climate-related SNPs by method. The horizontal bars show the number of candidate SNPs identified by each GEA method, while the vertical bars indicate the number of candidate SNPs unique to a single method and those shared across multiple methods, with the corresponding method(s) represented by the dots directly below the vertical bars. Colors refer to whether the method corrects (in green) or not (in orange) for population structure.

**Table 2.**
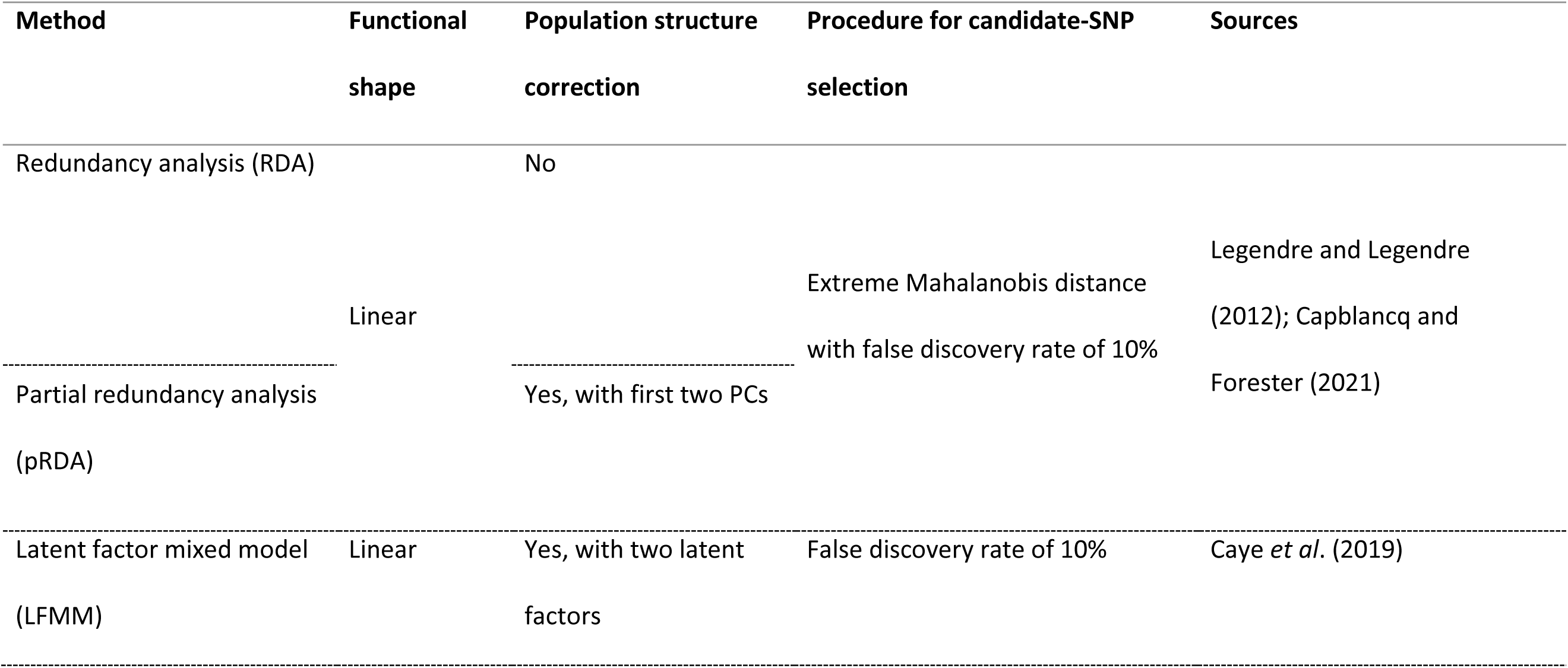
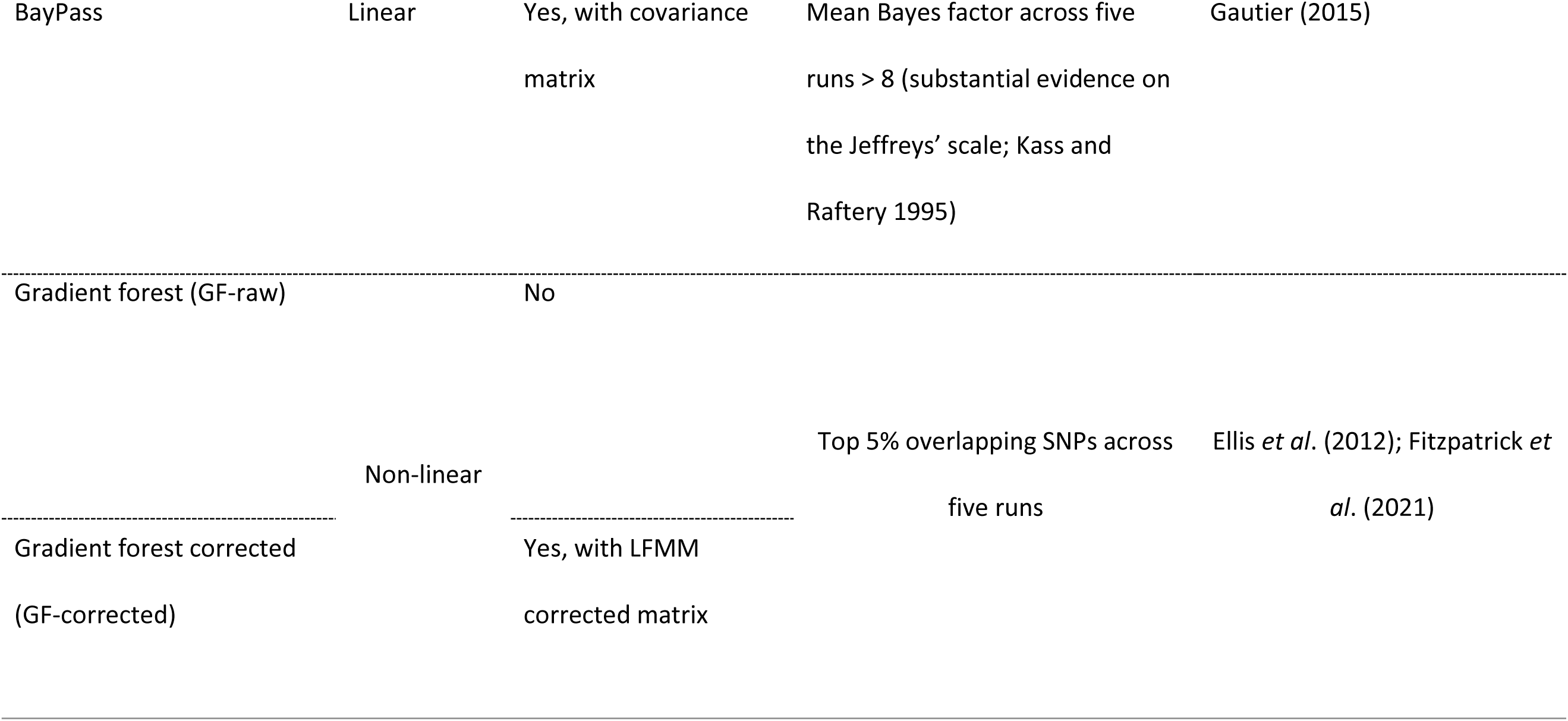
Description of the six genotype-environment association (GEA) analyses used to detect climate-associated loci (see additional details in Supplementary Material).

Candidate SNPs identified by at least two GEA methods were retained in an ‘*outlier set*’. This set was filtered to include only unlinked SNPs within contigs, using a linkage disequilibrium threshold of *R*² < 0.7, calculated using the genetics R package (Warnes 2022). A set of putatively neutral SNPs (hereafter ‘r*andom set*’) was generated by randomly selecting the same number of SNPs as in the *outlier set* from the SNPs that were not identified as candidates by any of the GEA methods, matching the allele frequencies of the *outlier set*. Finally, we also performed the downstream analyses using all the available SNPs (hereafter the ‘*all set*’).

#### Identifying populations deviating from GEA patterns

We identified populations that deviate from the global GEA trends provided by the RDA model with the best correlation with fitness proxies (i.e., the RDA model based on the *random set* of SNPs; see Figure 3). To achieve this, we extracted the observed and predicted population scores (using the reference time period 1901-1950) based on the linear combination of the climatic variables in the constrained RDA ordination space. These scores were extracted for the most explanatory RDA axes only (*K* = 2 in our case) and were weighted by axis importance. Euclidean distances between the observed and predicted population scores were calculated and subsequently standardised using a min-max normalization to range from zero to one. The resulting index is hereafter referred to as ‘genomic discrepancy index’ (GDI-RDA). Populations with higher GDI-RDA values deviate more from the mean GEA relationship.

**Figure 3.**
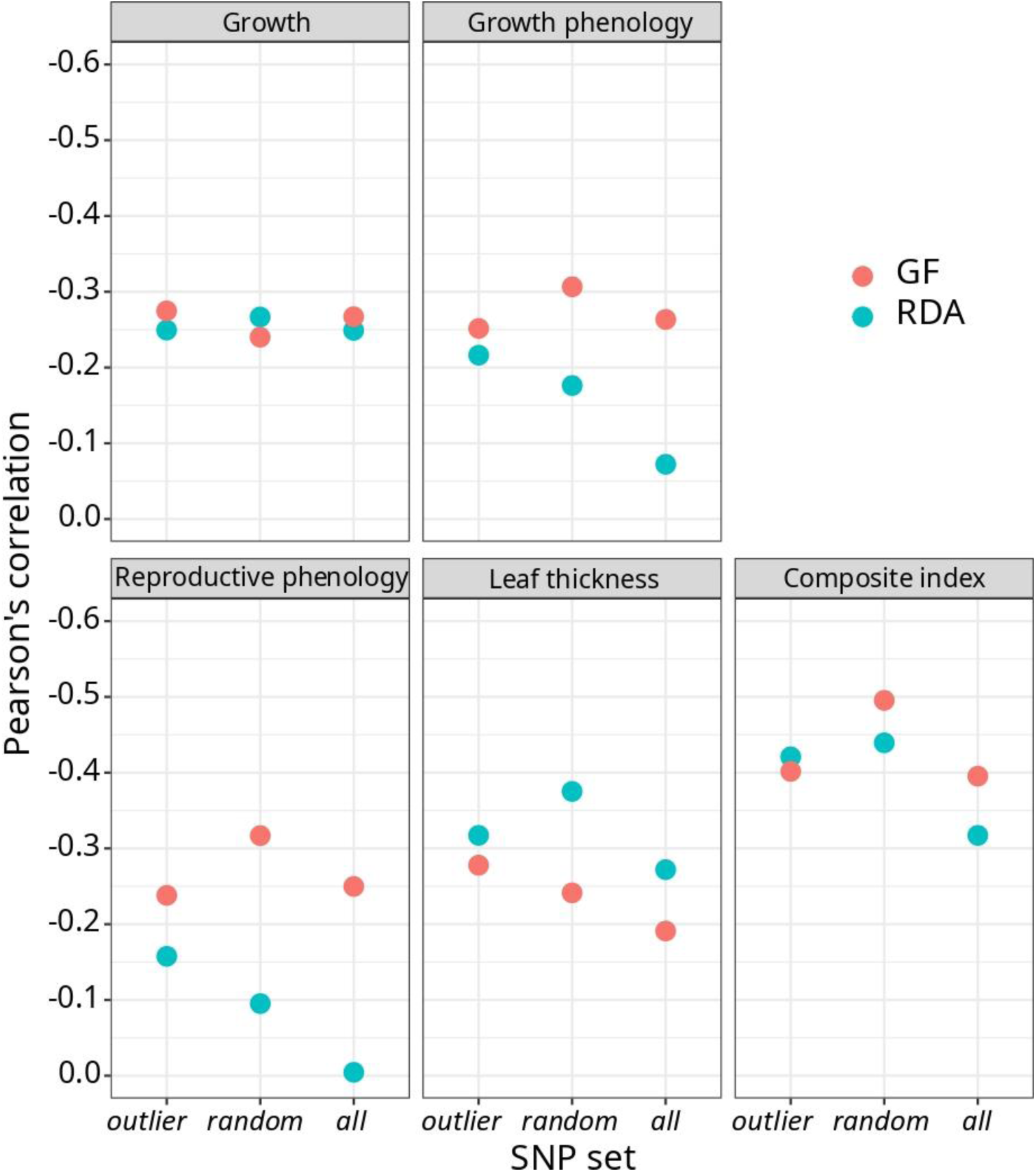
Pearson’s correlations between the genomic offset predictions and the fitness components assessed under common garden conditions. Genomic offset was calculated for each population as the Euclidean distance between the predicted optimal genomic composition in the climate of origin of the population for the reference period (1901-1950) and the predicted optimal genomic composition in the climate of the comparative experiment from the planting to the last measurement dates (1992-2012 for growth, growth phenology and reproductive phenology, and 1992-2021 for drought tolerance and the composite index), for each method and SNP set considered. The y-axis is inverted to display negative correlations in the upper part, reflecting our expectation that genomic offset predictions and phenotypic traits are negatively correlated under the specific climatic conditions of the comparative experiment (see main text and Figure S3).

### Potential maladaptation under future climate

#### Genomic offset predictions

The potential maladaptation of *T. baccata* populations to future climate was estimated using the genomic offset approach (Fitzpatrick and Keller 2015). Although several methods exist to compute this index, the general framework is similar. The first step estimates gene-climate relationships using genomic data and climate conditions at the population locations for a reference period (1901-1950 in our study). In the second step, these relationships are used to project the genomic composition of the populations for both the reference and future periods (2041-2070 under SSP3-7.0 in our study). The final step calculates the genetic distance between the reference and future projected genomic composition, i.e., the genomic offset. Genomic offsets were calculated using two methods, redundancy analysis (RDA; Capblancq and Forester 2021) and gradient forest (GF; Fitzpatrick and Keller 2015), whose main difference is the shape of the gene-climate relationship (i.e., linear for RDA and linear as well as nonlinear for GF). For both methods, we predicted the genomic offset using the three SNP sets described above (i.e., the *outlier set*, the *random set*, and the *all set*). We also used two additional SNP sets to assess the consistency of our predictions for the *random* and *outlier sets*: a second random SNP set, referred to as ‘*random_2*,’ and a set including all 935 outliers identified by at least one GEA method, referred to as ‘*all_outlier*’. All genomic offset predictions were made using five distinct GCMs (see Table S3) and standardised (min-max normalization). Genomic offset predictions were projected for a relatively near-future interval given the long generation time of *T. baccata* (reported to be as long as 75 years; Thomas and Polwart 2003), to address the assumption that genotype-climate associations based on current genomic data will remain valid for the prediction time period.

#### Evaluation of genomic offset predictions using phenotypic traits

To assess the relevance of genomic offset predictions, we estimated their Pearson’s correlations with the BLUPs of the four phenotypic traits (growth, growth phenology, reproductive phenology and drought/temperature tolerance) described above. Only populations with at least three individual plants measured for a given trait were included in the analyses. Single traits are often integrated into complex phenotypes, with different components of lifetime fitness being approximated by different traits, especially in long-lived organisms such as trees (see review in Climent *et al*. 2024). Therefore, we also correlated genomic offset predictions with a multi-trait index (hereafter ‘composite fitness index’), computed as follows:

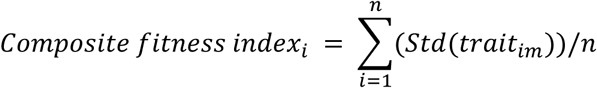

where *i* is the population, *trait_im_* denotes the population intercept (or BLUPs) of population *i* and trait *m*, *n* is the total number of traits and *Std* corresponds to a min-max standardisation (applied to each trait to constrain the values between 0 and 1). While simple, and with the lack of better information on trait-fitness correlation, this index combines all available traits and gives them equal weights.

Genomic offsets for populations from the comparative experiment were calculated using the GEA models built with the 29 SNP-genotyped populations and the Euclidean distance between the predicted optimal genomic composition at the population’s reference climate of origin (1901-1950 period) and the experimental site’s climate from the planting to the last measurement date (1992- 2012 for growth, growth phenology and reproductive phenology, and 1992-2021 for drought/temperature tolerance). Relationships between trait BLUPs and genomic offset predictions for each trait and the composite fitness index were investigated using both Pearson’s correlation coefficients, as suggested by Lotterhos (2024b), and linear models, as suggested by Fitzpatrick *et al*. (2021). For the latter, analyses were both conducted under a frequentist framework using the stats R package (R Core Team 2023) and under a Bayesian framework using the brms R package (Bürkner 2021; see Supplementary Material for details). Because of the particular climatic conditions of the comparative experiment, we expected a negative correlation between genomic offset predictions and phenotypic traits, assuming that later phenology, higher growth and larger leaf thickness would confer higher fitness in this environment.

## Results

### Climatic variation

Visual exploration of the climatic envelope of the 29 SNP-genotyped populations suggests that it adequately covers the climatic niche of *T. baccata*, as determined from the 298 sites with known presence of the species across its range (Figure S2a). The climatic envelope of the populations planted in the comparative experiment matched the climatic envelope of the 29 SNP-genotyped populations (Figure S2b). Finally, the climates of the source populations of the plants in the comparative experiment (reference period 1901-1950) were milder than the current climate of the experimental site (time periods 1992–2012 and 1992–2021), which is characterised by higher temperature, especially during the dry season, higher temperature seasonality, and lower annual precipitation (Figure S3). This showed that the experimental location was suitable for validating genomic offset predictions associated with climate change.

### Population genetic structure and historical gene flow

Bayesian clustering analysis identified two main gene pools (Western Europe and Eastern Europe) while centrally located populations, such as those from Switzerland and Germany, exhibited admixed genetic compositions. PCA revealed a similar geographic pattern, consistent with an east-west genetic structure (Figures S4–S6). According to *ΔK*, the best number of clusters was *K*=2 (Figure S6c), as largely confirmed by the topology of both individuals and populations along the first two PC axes (Figures S4 and S5). The Western Europe gene pool includes the populations from Spain, United Kingdom, Norway, France and southern Italy; while the Eastern Europe one includes those from Sweden, Germany, Switzerland, northern Italy, Slovenia, Slovakia, Bosnia-Herzegovina and Greece.

Population-specific *F*_ST_ ranged from 0.05 to 0.29, with most of the highest values found in the Mediterranean areas, but with no region consistently exhibiting the highest genetic differentiation (Figure 4a and Table S6). Additionally, EEMS models enabled identification of multiple populations with higher (e.g., Sainte-Baume in France, Omberg in Sweden, Gabrovo in Bulgaria or Tosande in Spain) or lower (e.g., Skarzynsko-Kamienna in Poland, Yenice in Turkey, Las Hurdes in Spain or the Middle Atlas in Morocco) migration than under an isolation by distance (IBD) model (Figure S7). Levels of estimated historical migration did not seem to follow any particular geographical or environmental pattern. In addition, the correlation between population-specific *F*_ST_ and historical migration estimates from EEMS for the 29 SNP-genotyped populations was very low and non-significant (Pearson’s correlation = −0.21, *p*-value = 0.27), suggesting that the species’ current genetic patterns likely result from multiple interacting processes (e.g., historical fragmentation, drift, irregular gene flow, or local adaptation).

**Figure 4.**
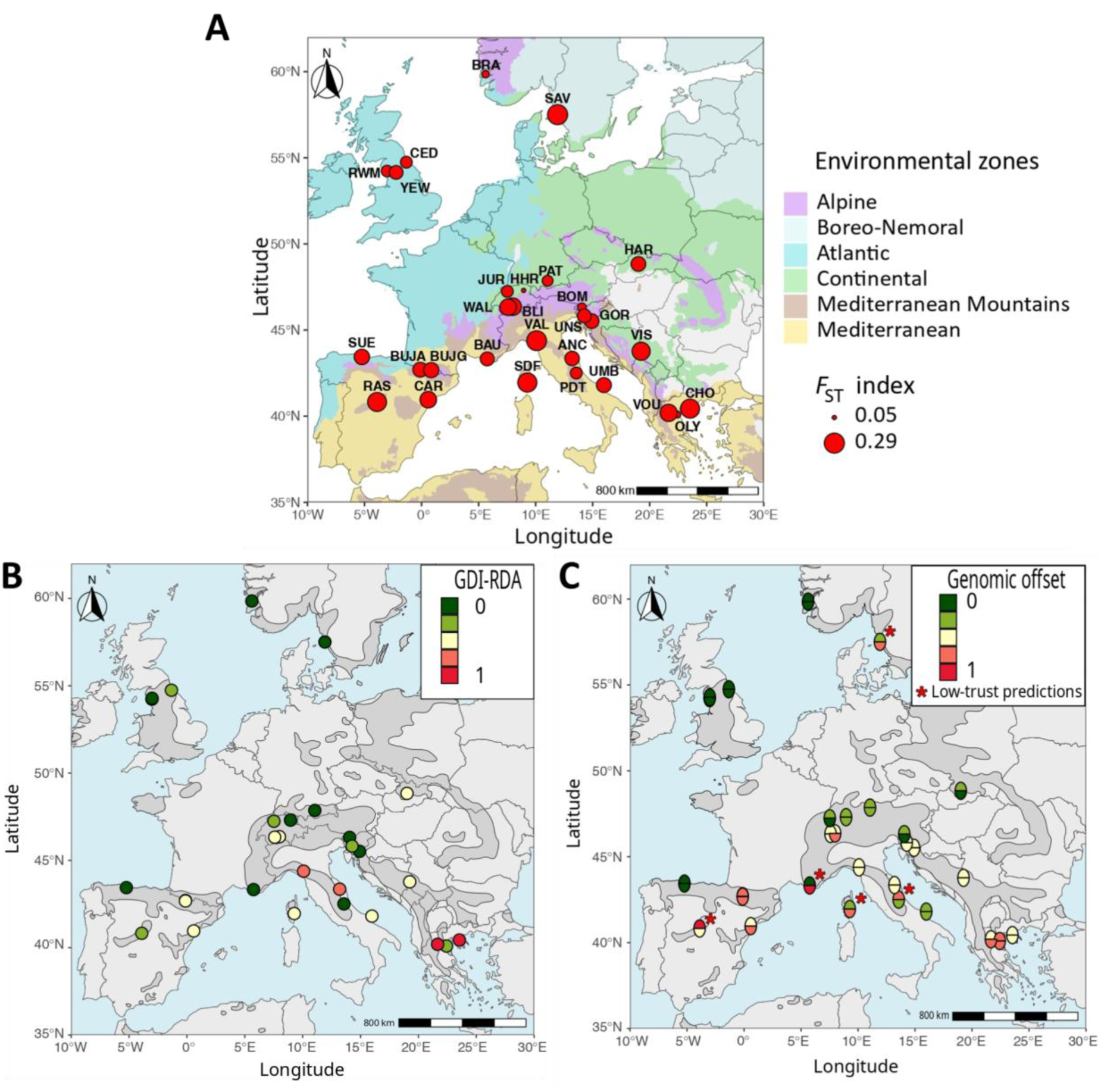
Potential maladaptation of *Taxus baccata* to future climate. A: population-specific *F*_ST_, an estimate of historically realised gene flow, on the European-scale environmental zone map of Metzger (2018). The size of the 29 red dots is proportional to their *F*_ST_ value. Population labels correspond to those in Table S1. B: Genomic discrepancy index (GDI-RDA) based on the *random set* of SNPs. A higher GDI-RDA indicates a greater difference between the predicted and observed genomic composition at present time. C: Genomic offset predictions for the RDA (lower half-circle) and GF (upper half-circle) models using the *random set* of SNPs and the mean climate across five general circulation models for the 2041-2070 time interval under the SSP3-7.0. Red stars represent the populations where the predictions of the two genomic offset models differed by more than two classes. In B and C, GDI-RDA and genomic offset predictions were standardized to the range 0-1 and categorised into five classes from low (green) to high (red) values. The species’ European range is presented in dark grey (EUFORGEN; Caudullo *et al*. 2017).

### Patterns of local adaptation to climate

#### Genetic variance partitioning

The full RDA model explained 58.6% of the total genetic variance across the 29 SNP-genotyped populations and provided strong evidence against the null hypothesis (i.e., no association between the model predictors and genetic variation; Table 3). Partial RDA models showed that 30.9% of the genetic variance was attributed solely to climate, 17.9% to the demographic history, and 22.2% to geography, while 29% of the genetic variance remained confounded between the predictors (see also Figures S5b and S8).

**Table 3.** Partition of the total amount of genetic variance showing proportions explained by climate (*clim*.), demographic history (*pop.struc*.) and geography (*geog*.), as well as by the confounding effect among these factors. See the main text for details.

| Partial RDA models | Proportion of total<br>explained variance ($R^2$) | Proportion of relative<br>explained variance | *p*-value |
| --- | --- | --- | --- |
| Full model: $F \sim clim. + pop.struc. + geog.$ | 0.586 | 1 | 0.001 |
| Pure climate: $F \sim clim. \mid (pop.struc. + geog.)$ | 0.181 | 0.309 | 0.041 |
| Pure population structure: $F \sim pop.struc. \mid (clim. + geog.)$ | 0.105 | 0.179 | 0.001 |
| Pure geography: $F \sim geog. \mid (clim. + pop.struc.)$ | 0.130 | 0.222 | 0.030 |
| Confounded climate/population structure/geography | 0.170 | 0.290 |  |
| Total unexplained | 0.414 |  |  |

#### Climate-related candidate SNPs

Genotype-environment association (GEA) methods identified different numbers of climate-related candidate SNPs (Figure 2 and Figures S9-S13 in Supplementary Material). RDA and pRDA identified the highest number of candidates (373 and 397 SNPs, respectively), and also had the largest number of candidates in common (57 SNPs). GF uncorrected for population genetic structure (GF-raw) and BayPass identified a similar number of candidate SNPs (110 and 133, respectively), while GF corrected for population genetic structure (GF-corrected) and LFMM identified the smallest number of candidate SNPs (59 and 24, respectively). The number of overlapping candidate SNPs between at least two methods was small compared to the overall number of candidate SNPs (136 vs. 935 SNPs). Furthermore, no candidate SNP was identified by more than three methods. After pruning SNPs for LD within contigs, 100 candidate SNPs were retained as outliers potentially involved in climate adaptation across populations (the ‘*outlier set*’).

#### Identifying populations deviating from GEA patterns

We found a general pattern of higher GDI-RDA at lower latitudes, supported by a negative Pearson’s correlation of −0.51 between GDI-RDA and latitude (*p*-value = 0.005), with Greek, Bosnian- Herzegovinian, Spanish and Italian populations having in most cases higher GDI-RDA than Swedish, Norwegian, British and Swiss ones (Figure 4b).

### Genomic offset predictions

#### Variability across methods and SNP sets

Variability in genomic offset predictions was observed across GCMs, with RDA predictions showing greater variability than those of GF (Figures S14 and S15). We used the mean genomic offset predictions across the five GCMs for subsequent comparisons and analyses (Table S8). For each method, genomic offset predictions across the main SNP sets were highly correlated (i.e., all Pearson’s correlations above 0.90 for RDA and 0.70 for GF; Figure S16) and thus population ranks based on genomic offset were largely consistent (Figure S17). Moreover, the *random_2* SNP set provided predictions consistent with the initial *random set*’s and the *all_outlier* SNP set also provided predictions very similar to the *outlier set*’s (all Pearson’s correlations above 0.90; Figure S18). In contrast, we observed some variability in genomic offset predictions across methods (albeit substantial correlation remained: lowest Pearson’s correlation = 0.30; see Figure S16). For instance, the Pearson’s correlation between RDA and GF using the *outlier set* was 0.41, resulting in discrepancies in population rankings (e.g., the southern French population of Sainte-Baume (BAU) had the highest genomic offset using RDA, but the second lowest using GF; Figure S17).

#### Evaluation using phenotypic traits

Evaluation of genomic offset predictions using phenotypic data provided consistent results for almost all genomic offset models. Indeed, according to Pearson’s correlations, all genomic offset predictions were negatively associated with the phenotypic traits (except for the RDA model based on *all set* and reproductive phenology), as expected, with significant *p*-values for several correlations (Figures 3 and S19, and Table S9). Several of these correlations were also supported by frequentist and Bayesian linear models. For instance, the negative correlations observed between RDA models (with *random* and *outlier sets*) and GF models (with *random set*) and the composite fitness index were highly unlikely to be obtained by chance (see Tables S10 and S11). Additionally, we found that the negative Pearson’s correlations between the genomic offset predictions and the composite fitness index were greater than most correlations for single traits (Figure 3). Finally, we also found that several genomic offset models appeared similarly associated with the composite index, and thus no prediction model can be confidently identified as displaying the best genomic offset predictions overall. Therefore, to account for the lack of a clearly identified best genomic offset model in the evaluation analyses, we retained the two models most strongly associated with fitness proxies and displayed both predictions on the same map for each population by organising predictions into five classes (Figure 4c).

Predictions were considered to be congruent (and therefore robust) only when the two models differed by less than two of these classes.

#### Geographical trends

Based on robust predictions (i.e., those supported by both GF and RDA models with *random set*), we found a negative correlation between genomic offset and latitude (Pearson’s correlation of −0.71, *p*- value = 0.0001), with most of the populations that had higher genomic offset being located at low latitudes in Mediterranean environments (VOU, OLY and CHO in Greece, CAR in Spain, and ANC and VAL in Italy). Additionally, we also found a positive correlation between genomic offset and elevation (Pearson’s correlation of 0.65, *p*-value = 0.0005). However, this correlation was substantially reduced when Atlantic low-elevation populations (CED, RWM and YEW in UK, and BRA in Norway) were removed (Pearson’s correlation of 0.39, *p*-value = 0.08), suggesting that the correlation is mostly driven by the very low genomic offset of Atlantic coastal populations (but notice the high genomic offset of some high-elevated populations: BLI in the Switzerland Alps and BUJA & BUJG in the Spanish Pyrenees). Finally, populations located in continental environments were also predicted to have low genomic offset (i.e., PAT in Germany or HAR in Slovakia). GF and RDA genomic offset models using other SNP sets (i.e., *outlier, random_2* and *all_outlier*) showed rather similar geographical patterns and comparable numbers of populations with inconsistent predictions (except for the models using the *outlier set*; see Figures S20-S22).

## Discussion

In this study, we found evidence of local adaptation to climate in *T. baccata*, by unveiling substantial genetic variation among populations explained by climate (18.1 % of the total genetic variance) and identifying a sizeable number of climate-associated loci using several GEA methods. We found variability in genomic offset predictions across methods (GF and RDA), but not across SNP sets (*all SNP, random, outlier*). Despite this, we found negative correlations between nearly all the genomic offset models and four phenotypic traits related to fitness (growth, growth phenology, reproductive phenology and drought tolerance) measured in plants from 26 nearly independent populations planted in a comparative experiment under common garden conditions. By combining information from estimates of historical genetic differentiation and effective migration, local adaptation and validated future climate maladaptation predictions, we provided valuable and credible insights into population vulnerability to future climate in *T. baccata*. Overall, our results highlight the vulnerability of Mediterranean and high-elevated mountain populations to ongoing climate change.

### Range-wide population genetic structure and historical gene flow

The two main gene pools identified at the European scale using SNP markers (Western and Eastern) are consistent with those previously reported by Mayol *et al*. (2015) based on nuclear microsatellites at a similar spatial scale. This study also detected a weak but significant isolation-by-distance (IBD) pattern across the entire species distribution. Nevertheless, despite these large-scale patterns, many *T. baccata* populations appear to be strongly isolated. This is supported by the high population- specific *F_ST_* values observed in our study, and a large body of literature reporting strong *F_ST_* values at local or regional geographical scales (e.g., Dubreuil *et al*. 2010; González-Martínez *et al*. 2010; Maroso *et al*. 2021; Casier *et al*. 2024; Chybicki *et al*. 2024). Notably, population-specific *F*_ST_ and EEMS provided divergent estimates of historical gene flow in many cases, likely reflecting the influence of multiple demographic processes beyond simple IBD. However, both methods converged in specific cases, e.g. VAL (Northern Italy) and CHO (Greece), which were consistently inferred to have experienced reduced historical gene flow, while HAR (Slovakia) and CED (United Kingdom) were among the populations with signals of historically high connectivity.

### Evidence supporting local adaptation to climate

Genotype-environment association (GEA) and the related genomic offset framework assume that populations are locally adapted (Rellstab *et al*. 2015). The overall genetic variance explained by climate predictors in *T. baccata* (18.1 %) was similar to or even higher than in other trees (*Fagus sylvatica* L.: 17 %, Capblancq *et al*. 2020b; *Pseudotaxus chienii* W.C. Cheng: 8.2 %, Liu *et al*. 2021; *Picea rubens* Sarg.: 14 %, Capblancq *et al*. 2023). In addition, we identified a substantial number of climate-related candidate SNPs in this species, including two from genes previously reported to be potentially under positive selection for climate in *T. baccata* (Table S12; Mayol *et al*. 2020).

Nevertheless, congruent genomic offset predictions and correlations with phenotypic traits across SNP sets (including random sets) suggest that the signals captured by GEA may result from linkage disequilibrium (LD) with genome-wide loci of small effect, and thus that climate adaptation along the identified climate gradients is polygenic rather than oligogenic in this species. This hypothesis is supported by recent literature on trees, which report widespread polygenic adaptation for adaptive traits (e.g., De La Torre *et al*. 2019; de Miguel *et al*. 2022).

One of the main assumptions underlying GEA and genomic offset methods is that populations are equally locally adapted to their environments (Ahrens *et al*. 2023; Lotterhos 2024b). However, numerous studies in diverse species have shown that this assumption is often not met. For example, Marcora *et al*. (2021) found higher performance (e.g., germination success, sapling growth and survival) under site-of-origin conditions in *Polylepis australis* (Bitter), an endemic tree species of central Argentina, but only in marginal populations. There is also evidence of adaptation lags to current climate affecting unequally different parts of a species’ range, probably associated with contrasting demographic histories (Browne *et al*. 2019; Fréjaville *et al*. 2020). Furthermore, in presence of complex mechanisms such as conditional neutrality or genetic redundancy, GEA may fail to detect adaptive loci (Rellstab *et al*. 2015; Lind *et al*. 2018). To assess variability in local adaptation across populations or potential differences in adaptive trajectories, we calculated a new ‘genomic discrepancy index’ (GDI-RDA, see Methods). Interestingly, in this study, GDI-RDA followed the inverse pattern of historical effective migration detected by EEMS (Pearson’s correlation of −0.39, *p*-value = 0.03), with higher GDI-RDA associated with lower historical migration rates. This pattern was also supported by population-specific *F*_ST_ (Pearson’s correlation of 0.52, *p*-value = 0.004). This suggests that isolation may have hindered local adaptation or led to different adaptive trajectories by limiting the influx of beneficial alleles other populations (e.g., Deacon and Cavender-Bares 2015) or, alternatively, may have exacerbated the effects of genetic drift (Lande 1993). However, inconsistencies between estimates derived from EEMS and population-specific *F*_ST_ highlight the need for further research to clarify the role of gene flow in local adaptation in *T. baccata*. Here, we used GDI-RDA to assess the degree of local adaptation by identifying populations deviating from the overall GEA patterns identified by RDA. To test empirically the validity of this index, future work could use reciprocal transplant experiments (Kawecki and Ebert 2004). The discrepancy index approach could also be extended beyond RDA to other GEA methods (e.g., gradient forest, which does not assume a linear relationship) or incorporating this information directly into the genomic offset framework.

### Towards a more comprehensive genomic offset framework

We found notable variability in genomic offset predictions across methods (GF and RDA). However, despite these differences, we still found substantial positive correlations among them, regardless of the SNP set used, and mostly consistent population rankings. These results are reassuring compared to those of Archambeau *et al*. (2025), who found almost no correlations between RDA and GF predictions in maritime pine (*Pinus pinaster* Aiton), another European conifer, and inconsistent population rankings. Similarly, Lind *et al*. (2024a) reported inconsistent genomic offset predictions between GF and the Risk of Non-Adaptiveness (RONA, Rellstab *et al*. 2016) for Jack pine (*Pinus banksiana* Lamb.), but not for Douglas-fir (*Pseudotsuga menziesii* Mirb.). These stark differences between studies illustrate a likely species-specific effect on the consistency of genomic offset predictions across methods, possibly related to levels of population genetic structure, shape of climate adaptive clines and/or the genomic sampling used (e.g., candidate genes vs. random markers). Taken together, these studies highlight that, in the current framework, the use of genomic offset predictions without empirical validation should be avoided.

Empirical validation in our study was performed by using phenotypic traits measured under common garden conditions. We found that random SNP sets had predictive accuracy similar to that of candidate SNPs identified by GEAs. This is in agreement with recent studies on genomic offset prediction based on GF (Fitzpatrick *et al*. 2021; Lind *et al*. 2024a) and RDA (Archambeau *et al*. 2025), which have also reported comparable performance between candidate and random marker sets. In our study, these results might be explained by the fact that both the candidate and random SNPs were drawn from coding regions. As a result, even the random SNPs may carry biologically relevant signals due to linkage with functional variants, as also noted by Lind *et al*. (2024a). Moreover, as the phenotypic traits used for validation are expected to be highly polygenic (see, e.g., de Miguel *et al*. 2022 for growth), predictive models may benefit from diffuse genome-wide LD patterns rather than individual large-effect SNPs.

Almost all the genomic offset models had moderate to substantial negative correlations with the phenotypic traits, consistent with current knowledge on the adaptation of trees to Mediterranean mountains (Liu *et al*. 2020; Arroyo *et al*. 2021; Quan *et al*. 2024). While a negative relationship between genomic offset predictions and fitness proxies has already been demonstrated using common gardens (Rhoné *et al*. 2020; Fitzpatrick *et al*. 2021; Lachmuth *et al*. 2023; Archambeau *et al*. 2025), to our knowledge, it has never been tested before using fitness proxies from a nearly independent set of populations. Therefore, our study addresses a pending question in the literature (Lotterhos 2024b), by suggesting that the observed negative correlation between genomic offset predictions and fitness under common garden conditions is not the result of model overfitting.

Finally, we found a stronger negative association between genomic offset and the index integrating the different phenotypic traits (i.e., the composite fitness index) than for any single trait. This finding suggests (1) that phenology and drought/temperature tolerance traits may provide information on fitness components other than growth and, thus, (2) that combining multiple traits into a synthetic index may help to better approximate fitness, especially for long-lived organisms, such as trees, for which direct lifetime fitness cannot be measured (Climent *et al*. 2024). Nevertheless, although promising, the calculation of the composite index in this study is rather simplistic and may not properly account for the complex interaction of phenotypic traits that determines fitness, and calls for further work on methods for phenotypic trait integration in trees.

To conclude, we argue that studies assessing population maladaptation to climate in a conservation context should restrict their estimates to the genotyped populations only (i.e., to adopt a more ‘population-centred’ framework), and only when predictions from different methods are in agreement. Indeed, extrapolating genomic offset models to unsampled areas can lead to misleading conclusions in the context of population conservation (see Lind and Lotterhos 2024b). We have also shown in this study that conflicting genomic offset predictions can arise across models for some genotyped populations, even when the predictions are linked with fitness proxies using empirical data. Moreover, integrating additional information, such as the degree of local adaptation or connectivity, seems advisable for a more comprehensive assessment of population vulnerability in the face of climate change.

### The fate of English yew in the face of climate change

Given the long-term demographic decline in *T. baccata*, due to various factors including long-term climate variability, one may expect climate change to be an additional threat for population survival (Bach *et al*. in revision). By combining information on the potential degree of local adaptation (GDI- RDA), historical genetic differentiation, effective migration surfaces, and potential maladaptation to future climate, we identified several *T. baccata* populations potentially at risk in the face of climate change.

Populations located along the Mediterranean coast and in high-elevation mountain environments (i.e., in Greece, northern and central Italy, Pyrenees, Mediterranean Spain, and southern Switzerland) mostly shared similar features regarding the above-mentioned factors, suggesting a higher vulnerability to future climate. Apart from high genomic offsets, it appears that most of these populations have evolved under strong historical isolation, so their access to beneficial alleles through gene flow may be also compromised in the near future if these patterns are maintained. In addition, most of these populations showed substantial GDI-RDA, suggesting that they deviate from average climate adaptation patterns in the species. This may be because of pre-adaptation, distinct adaptive patterns not captured by the statistical framework, or a lower degree of adaptation due to adaptation lags or random genetic drift. In the latter case, the expected maladaptation to future climate of high GDI-RDA populations may be even worse than estimated by genomic offsets.

Overall, our results are in accordance with the current body of literature that shows the considerable impact of climate change on tree species inhabiting Mediterranean and high-elevation mountain regions (Capblancq *et al*. 2020; Calama *et al*. 2024; Longo-Minnolo *et al*. 2025). They also suggest that *T. baccata* populations in Atlantic and continental environments may cope better with future climate. Indeed, populations from northern Spain, United Kingdom, Norway, Germany and Slovakia have very low genomic offsets, follow overall climate adaptation patterns (i.e., low GDI-RDA) and, in most cases, seem to have evolved under lower historical isolation.

However, caution must be exercised while considering these interpretations, as there are several limitations to our analyses. First, population-specific *F*_ST_ and EEMS provided inconsistent information on the historical gene flow patterns suggesting a complex demographic history that is not fully accounted for by either of the two approaches. Additionally, these analyses provided information on historical patterns but not on the current or future gene flow patterns, which remain unknown. At present, many *T. baccata* populations have low population size or suffer from habitat fragmentation, and thus may even be more isolated than in the past, as shown in other tree species with currently fragmented distributions (Jump and Peñuelas 2006; Sebbenn 2011). Second, the genomic offset framework relies on several assumptions that may limit its validity (see Rellstab *et al*. 2021; Ahrens *et al*. 2023, amongst others). For example, these approaches assume that the genotype-environment relationships will remain constant, thus neglecting the adaptive capacity of populations. We have addressed some of these assumptions by using common garden data to establish a relationship between genomic offset predictions and fitness, and by limiting our predictions to near-future time intervals, making standing genetic variation the most likely source of future adaptation (Barrett and Schluter 2008). Nevertheless, we must remain aware of other untested assumptions that may impact future climate maladaptation predictions (e.g., overall lack of local adaptation). Furthermore, the insights into future climate vulnerability provided in this study do not take into account other processes that may mitigate the effects of climate change in the short term, such as phenotypic plasticity, or exacerbate its effects, such as inbreeding depression (Capblancq *et al*. 2020a; Aguirre-Liguori *et al*. 2021). More broadly, biotic interactions, such grazing intensification, pest outbreaks and competition, may also be altered by climate change and have a strong impact on *T. baccata* populations (Thomas and Polwart 2003; Mysterud and Østbye 2004; Ruprecht *et al*. 2010).

## Conclusions

Our study demonstrates the complexity of applying the genomic offset framework for the genetic conservation of *T. baccata*, a declining tree species. It also highlights the benefits of combining different sources of information (e.g., genomic and phenotypic data) and analyses (e.g., genomic offsets, local adaptation estimates and historical gene flow), to gain insight into population vulnerability to climate change. By using a nearly independent set of populations growing under common garden conditions, we provided a robust interpretation of genomic offset predictions and showed how integrating multiple phenotypic traits in simple indices can help to approximate fitness for long-lived organisms such as trees. However, we also found potential variability in the degree of local adaptation across populations and different degrees of historical isolation, which were not fully consistent across estimation methods. Finally, by combining insights into levels of local adaptation and historical effective gene flow with maladaptation predictions, we suggest that European Mediterranean and high-elevation mountain areas populations of *T. baccata* will suffer the most in the face of climate change, as opposed to Atlantic and continental ones.

## Supporting information

none

## Acknowledgements

We wish to thank Thibaut Capblancq for helpful insights and discussion, and Natalija Dovč, Tjaša Baloh, Rok Damjanić, Carlo Urbinati, Alessandro Vitali, Camilla Avanzi, Ilaria Spanu, Celeste Labriola, Daniele Di Santo, Paolo Piovani, Simone Barbarotti, Stefano Zanzucchi, Eleftheria Dalmaris, and the personnel of Reparto Carabinieri Biodiversità Foresta Umbra for field work. TF was funded by the INRAE scientific department ECODIV. EV was supported by the MSCA European fellowship MedForAct (GA 101107604). FAA was supported by project Taxol-GR (MIS5004922) co-financed by Greece and the EU. Sampling in Slovenia and MW were funded by Slovenian Research Agency (research core funding no. P4-0107). This research was funded also by EU H2020 GENTREE (grant agreement No. 676876), TAXUS (CGL2007-63107/BOS), ADAPCON (CGL2011-30182-C02-02), READAPT (PID2020-112738GB-I00), and EU H2020 FORGENIUS (grant agreement No. 862221) projects. Views and opinions expressed are however those of the authors only and do not necessarily reflect those of the European Union. Neither the European Union nor the granting authority can be held responsible for them.

## Data Archiving Statement

Genomic data are openly available in Data INRAE at DOI: 10.57745/JJ0ZEI. Scripts for genotype-environment association (GEA), and genomic offset computation and validation can be found in a GitHub public repository at https://github.com/Thomas-Francisco/Genomic-signatures-of-mal-adaptation-to-climate-in-English-Yew.

## Author Contributions

Designed research: SCG-M, TF; field work: MM, MR, MW, FB, GGV, AP, SC; genetic data acquisition and pretreatment: SP, SCG-M, FB, GGV, MM, MR, TF; phenotypic data acquisition and pretreatment: MR, MM; data analysis: TF, EV; funding: MM, MR; wrote the paper: TF, SCG-M. All authors critically read, edited, commented and approved the submitted version of the paper.

## Notes

### Competing Interest Statement

The authors have declared no competing interest.

