## Supplementary material for "Genomic signatures of climate-driven (mal)adaptation in an iconic conifer, the English yew (*Taxus baccata* L.)": none

### Section A: Population sets, climatic and phenotypic data

*Table S1.* Information on the 29 SNP-genotyped populations. *N* corresponds to the number of sampled trees per population (before SNP-quality filtering). Elevation data have been extracted from the Copernicus Digital Surface Model at a 90-metre resolution.

| Country | Population | ID | *N* | Latitude (°) | Longitude (°) | Elevation (m) |
|---|---|---|---|---|---|---|
| Bosnia-Herzegovina | Višegrad | VIS | 10 | 43.77 | 19.25 | 824 |
| France | Sainte Baume | BAU | 17 | 43.33 | 5.75 | 748 |
| France | Serra di Fiumorbu (Corsica) | SDF | 19 | 41.95 | 9.27 | 826 |
| Germany | Paterzeller Eibenwald | PAT | 18 | 47.86 | 11.05 | 647 |
| Greece | Mount Cholomon | CHO | 16 | 40.44 | 23.54 | 655 |
| Greece | Mount Vourinos | VOU | 19 | 40.19 | 21.66 | 1226 |
| Greece | Mount Olympus | OLY | 18 | 40.09 | 22.43 | 816 |
| Italy | Ancona | ANC | 27 | 43.35 | 13.20 | 668 |
| Italy | Prati di Tivo | PDT | 19 | 42.51 | 13.57 | 1510 |
| Italy | Foresta Umbra | UMB | 19 | 41.81 | 15.98 | 761 |
| Italy | Valditacca | VAL | 19 | 44.38 | 10.08 | 1074 |
| Norway | Brandvik Stord Hordaland | BRA | 18 | 59.85 | 5.61 | 31 |
| Slovakia | Harmanec | HAR | 10 | 48.84 | 19.01 | 812 |
| Slovenia | Gorenja žaga Kotnica | GOR | 19 | 45.52 | 14.91 | 223 |
| Slovenia | Lake Bohinj Obrne Mokri | BOM | 32 | 46.31 | 14.06 | 667 |
| Slovenia | Unška Koliševka | UNS | 19 | 45.82 | 14.27 | 626 |
| Spain | Bujaruelo ADAPCON | BUJA | 29 | 42.70 | -0.12 | 1388 |
| Spain | Bujaruelo GENTREE | BUJG | 19 | 42.67 | -0.12 | 1246 |
| Spain | Cardó | CAR | 10 | 40.96 | 0.59 | 619 |
| Spain | Rascafría | RAS | 10 | 40.82 | -3.90 | 1603 |
| Spain | Sueve | SUE | 10 | 43.44 | -5.23 | 788 |
| Sweden | Särö Västerskog | SAV | 18 | 57.51 | 11.92 | 22 |
| Switzerland | Blindtal | BLI | 10 | 46.36 | 7.99 | 1239 |
| Switzerland | Hörnli-Hulftegg Roten | HHR | 19 | 47.30 | 8.95 | 956 |
| Switzerland | Jura | JUR | 10 | 47.24 | 7.52 | 715 |
| Switzerland | Wallis | WAL | 10 | 46.33 | 7.59 | 1093 |
| United Kingdom | Castle Eden Dene | CED | 10 | 54.74 | -1.33 | 77 |
| United Kingdom | Roudsea Woods and Mosses | RWM | 26 | 54.23 | -3.02 | 18 |
| United Kingdom | Yewbarrow | YEW | 10 | 54.28 | -3.01 | 56 |

*Table S2*. Summary of the main genetic analyses conducted in this study and their associated datasets. MAC: minor allele count; nuSSRs: nuclear microsatellites

| **Type of analysis** | **Method** | **Population dataset** | **Genomic dataset** | **Climatic period** |
|---|---|---|---|---|
| Population genetic structure | Principal component analysis (PCA)<br>Bayesian clustering analysis (STRUCTURE) | 29 populations with genomic data | 8,252 SNPs not filtered by MAC | Not used |
| $F_{ST}$ index | Population specific divergence index using Bayescan V2.1 | 29 populations with genomic data | 8,252 SNPs not filtered by MAC | Not used |
| Historical effective gene flow | Estimated effective migration surfaces (EEMS) | 176 populations with nuclear microsatellite (nuSSR) data | 7 nuSSRs | Not used |
| Variance partitioning | Redundancy analysis (RDA) and partial redundancy analysis (pRDA) | 29 populations with genomic data | 8,616 SNPs filtered by MAC | Not used |
| Genotype-environmen analysis (GEA) | RDA<br><br>pRDA<br>Latent factor mixed model (LFMM)<br>BayPass<br>Gradient forest (GF-raw)<br>Gradient forest corrected for population structure (GF-corrected) | 29 populations with genomic data | 8,616 SNPs filtered by MAC | 1901-1950 reference period |

| | | | | |
|---|---|---|---|---|
| Genomic discrepancy index (GDI-RDA) | RDA | 29 populations with genomic data | 8,616 SNPs filtered by MAC | 1901-1950 reference period |
| Genomic offset | Gradient forest<br>RDA | 29 populations with genomic data | 8,616 SNPs filtered by MAC | 1901-1950 for reference period and 2041-2070 for future period |
| Genomic offset comparative experiment | Models built in the genomic offset section above were used to project the genomic composition of the 26 populations grown in the comparative experiment | | | 1901-1950 for population-origin reference climate and 1992-2012 or 1992-2021 for comparative experiment climate depending on the fitness proxy considered |

*Table S3*. The five global climate models (GCMs) used to predict the genomic offset under the socio-economic pathway 3-7.0 for the 2041-2070 period.

| **Global climate models (GCMs)** | **Source**[1] |
|---|---|
| GFDL-ESM4.1 | Dunne *et al*. (2020) |
| IPSL-CM6A-LR | Boucher *et al*. (2020) |
| MPI-ESM1-2-HR | Gutjahr *et al*. (2019) |
| MRI-ESM2-0 | Yukimoto *et al*. (2019) |
| UKESM1-0-LL | Sellar *et al*. (2019) |

[1]References :

Boucher O, Servonnat J, Albright AL, *et al.* (2020). Presentation and Evaluation of the IPSL-CM6A-LR Climate Model. Journal of Advances in Modeling Earth Systems 12:e2019MS002010.

Dunne JP, Horowitz LW, Adcroft AJ, *et al.* (2020). The GFDL Earth System Model Version 4.1 (GFDL-ESM 4.1): Overall Coupled Model Description and Simulation Characteristics. Journal of Advances in Modeling Earth Systems 12:e2019MS002015.

Gutjahr O, Putrasahan D, Lohmann K, *et al.* (2019). Max Planck Institute Earth System Model (MPI-ESM1.2) for the High-Resolution Model Intercomparison Project (HighResMIP). Geoscientific Model Development 12:3241–3281.

Sellar AA, Jones CG, Mulcahy JP, *et al.* (2019). UKESM1: Description and Evaluation of the U.K. Earth System Model. Journal of Advances in Modeling Earth Systems 11:4513–4558.

Yukimoto S, Kawai H, Koshiro T, *et al.* (2019). The Meteorological Research Institute Earth System Model Version 2.0, MRI-ESM2.0: Description and Basic Evaluation of the Physical Component. Journal of the Meteorological Society of Japan. Ser. II 97:931–965.

*Table S4*. Best climatic predictors to explain genomic variation across populations based on 20 *ordiR2step* runs.

| **Variable** | **Run count** |
|---|---|
| Bio 2: Mean Diurnal Range (°C) | 20 |
| Bio 4: Temperature Seasonality (standard deviation x100) | 20 |
| Bio 7 : Temperature Annual Range (°C) | 20 |
| Bio 9: Mean Temperature Driest Quarter (°C) | 20 |

*Table S5*. Phenotypic traits used as fitness proxies and the number of populations where at least three individuals were measured in the comparative experiment. The Gelman-Rubin criteria of the Best Linear Unbiased Predictors (BLUPs) calculation is also shown. Additional details on the measurement and calculation of these phenotypic traits are provided in the main text.

| **Phenotypic trait** | **Type** | **Number of populations** | **Gelman-Rubin criteria** |
|---|---|---|---|
| Shoot volume ($mm^3$) | Growth | 24 | 1.05 |
| Shoot elongation (proportion) | Growth phenology | 22 | 1.02 |
| Proportion open male strobili (proportion) | Reproductive phenology | 14 | 1.01 |
| Leaf thickness (mm) | Drought/temperature tolerance | 23 | 1.02 |

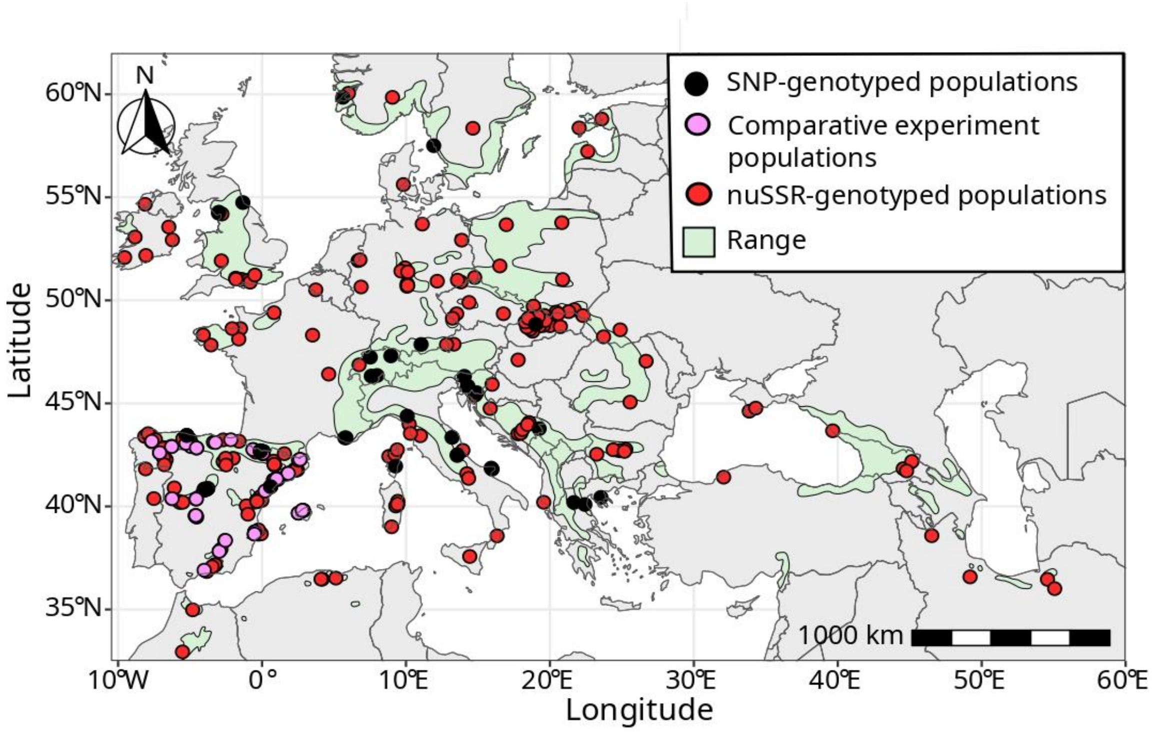


*Figure S1*. Map showing the three sets of *Taxus baccata* populations used in this study. The black dots represent the 29 SNP-genotyped populations used for the genomic analyses, the pink dots indicate the 26 independent populations planted in the comparative experiment under common garden conditions in central Spain and the red dots represent the 243 populations sampled for nuclear microsatellite (nuSSR) genotyping.

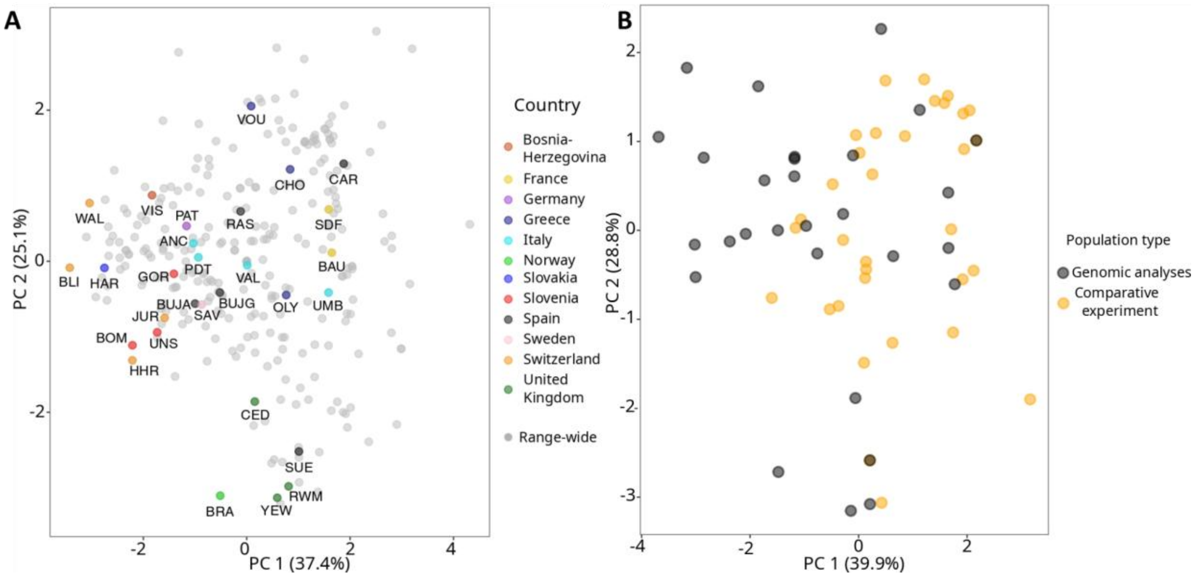


*Figure S2*. Principal component analysis (PCA) showing the climatic envelope of A: 29 populations used for genomic analysis coloured by country and the 298 sampled sites used in this study (grey dots, referred to as “Range-wide” in the legend), which represents the full range of climatic conditions where the species occurs in Europe; and B: 29 populations used for genomic analyses (black dots) and 26 populations planted in a comparative experiment under common garden conditions (orange dots).

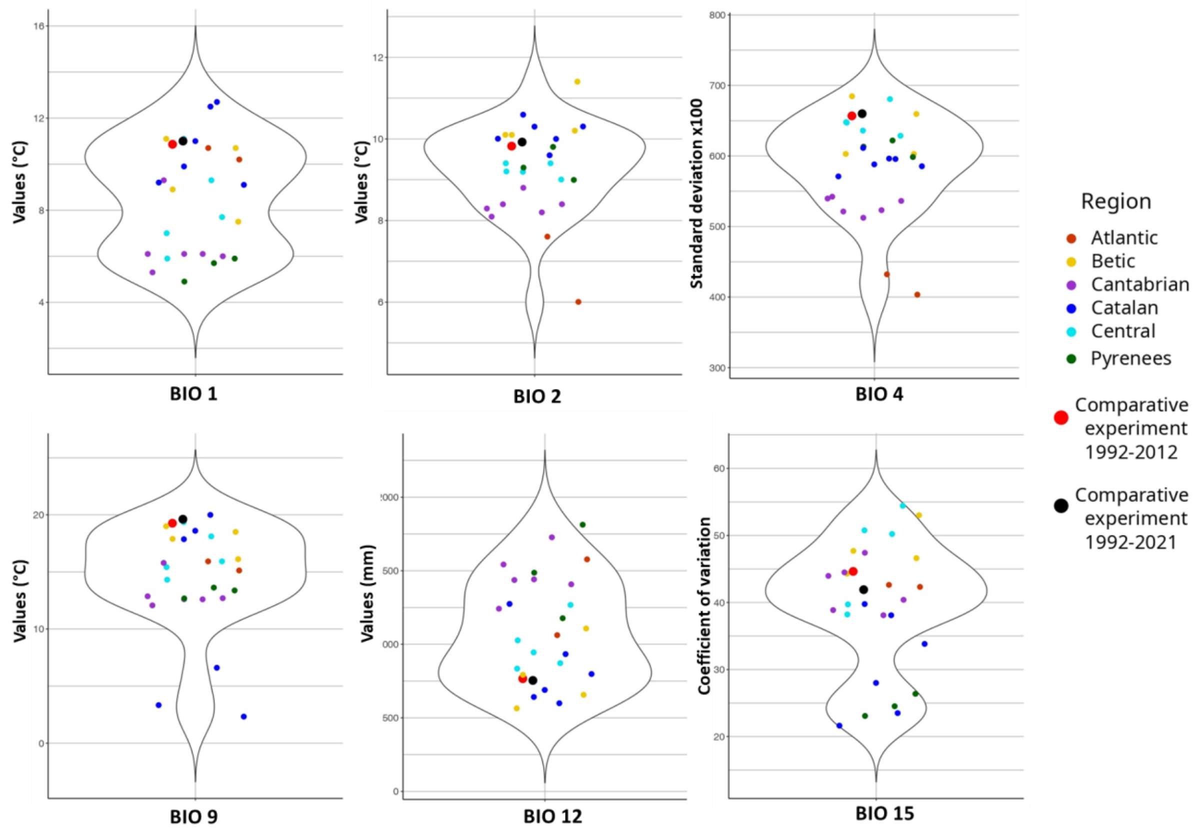


*Figure S3*. Violin plots displaying the climate conditions for the six retained climatic predictors of the 26 populations planted in the comparative experiment (1901-1950 time interval) as well as the climatic conditions of the comparative experiment for the two periods of trait measurements (see legend). Bio 1: Mean Annual Temperature; Bio 2: Mean Diurnal Range; Bio 4: Temperature Seasonality; Bio 9: Mean Temperature Driest Quarter; Bio 12: Mean Annual Precipitation and Bio 15: Precipitation Seasonality.

### Section B: Population genetic structure and historical gene flow

*Additional methods: details on EEMS computations*

Populations with potential historical permeability or barriers to gene flow across the range of *T. baccata* were identified by estimating effective migration surfaces (EEMS, Petkova *et al*. 2016). We tested different numbers of predicted demes (i.e., 100, 400, and 1,000 demes) as recommended by Petkova *et al*. (2016). For each deme configuration, EEMS was run three times independently using 2,000,000 iterations, with a burn-in period of 1,000,000 iterations and a thinning interval of 9,999. Default parameterisation was used except where parameter acceptance rates were outside the recommended ranges (i.e., 10-40%), in which case input parameters were adjusted according to the manual instructions. For each run, we calculated the $R^2$ of the relationship between observed and fitted dissimilarities in order to assess model-fit quality and select the best-performing model. EEMS models predicted on the basis of 1,000 demes had a greater association between observed and fitted dissimilarities than models with fewer predicted demes (Table S7). We also found more consistent results across runs for the models using populations with at least ten trees. Therefore, these settings were preferred for the final runs (Figure S7).

*References*

Petkova D, Novembre J, Stephens M (2016) Visualizing spatial population structure with estimated effective migration surfaces. Nat Genet 48:94–100. https://doi.org/10.1038/ng.3464

*Table S6.* Population genetic structure results for the principal component analysis (scores for each population on PC1 and 2) and the STRUCTURE analysis (proportional assignment probabilities for each population to the two major gene pools), as well as historical gene flow results for the EEMS analysis (spatial variation in effective migration rates across the landscape), and pairwise $F_{ST}$ estimates (levels of genetic differentiation among populations) for the SNP-genotyped populations.

| Population | PC 1 | PC 2 | Western gene pool | Eastern gene pool | Historical gene flow (EEMS) | $F_{ST}$ index |
|---|---|---|---|---|---|---|
| VIS | 6.75 | 3.94 | 0.05 | 0.95 | -0.12 | 0.23 |
| BAU | -3.66 | -0.41 | 0.88 | 0.12 | 1.49 | 0.14 |
| SDF | -2.81 | 3.11 | 0.86 | 0.14 | -0.34 | 0.26 |
| PAT | 1.76 | -4.69 | 0.33 | 0.67 | 0.48 | 0.09 |
| CHO | 5.61 | 4.89 | 0.08 | 0.92 | -0.31 | 0.25 |
| VOU | 2.08 | 3.28 | 0.07 | 0.93 | -0.73 | 0.21 |
| OLY | 5.32 | 3.53 | 0.43 | 0.57 | 0.71 | 0.05 |
| ANC | -2.21 | 3.04 | 0.83 | 0.17 | 0.04 | 0.15 |
| PDT | -1.39 | 3.67 | 0.73 | 0.27 | 0.16 | 0.10 |
| UMB | -0.74 | 4.07 | 0.68 | 0.32 | -0.05 | 0.16 |
| VAL | 2.54 | -3.38 | 0.21 | 0.79 | -0.29 | 0.28 |
| BRA | -0.47 | -0.27 | 0.57 | 0.43 | -0.76 | 0.06 |
| HAR | 8.36 | 2.21 | 0.02 | 0.98 | 0.20 | 0.15 |
| GOR | 4.49 | 0.02 | 0.13 | 0.87 | 0.15 | 0.15 |
| BOM | 4.78 | 1.27 | 0.16 | 0.84 | 0.43 | 0.07 |
| UNS | 5.58 | 0.80 | 0.08 | 0.92 | 0.15 | 0.13 |
| BUJA | -5.20 | 2.37 | 0.99 | 0.01 | -0.38 | 0.15 |
| BUJG | -5.07 | 2.81 | 1.00 | 0.00 | -0.38 | 0.16 |
| CAR | -5.50 | 1.97 | 1.00 | 0.00 | 0.83 | 0.19 |
| RAS | -5.82 | 3.42 | 1.00 | 0.00 | 0.01 | 0.25 |
| SUE | -5.41 | 1.43 | 0.98 | 0.02 | 0.95 | 0.17 |
| SAV | 1.78 | -3.48 | 0.29 | 0.71 | 0.57 | 0.29 |
| BLI | 0.61 | -10.91 | 0.34 | 0.66 | -0.48 | 0.22 |
| HHR | -0.71 | -4.94 | 0.53 | 0.47 | 0.96 | 0.05 |
| JUR | -0.14 | -6.58 | 0.46 | 0.54 | -0.54 | 0.10 |
| WAL | 0.13 | -9.85 | 0.39 | 0.61 | 0.39 | 0.19 |
| CED | -3.80 | -0.36 | 0.84 | 0.16 | 0.51 | 0.10 |
| RWM | -3.29 | -0.15 | 0.82 | 0.18 | -0.44 | 0.10 |
| YEW | -3.57 | -0.82 | 0.84 | 0.16 | -0.44 | 0.13 |

*Table S7*. Comparison of the estimated effective migration surfaces (EEMS) models carried out using two types of input data, three numbers of predicted demes and three replicates per model. For each model, the $R^2$ of observed to fitted dissimilarities was calculated to assess the quality of the model fit. Bold font indicates the model used in Figure S7.

| Input data | Number of predicted demes | Replicates | $R^2$ observed vs fitted dissimilarities |
|---|---|---|---|
| All populations (243 populations) | 100 | 1 | 0.322 |
| | | 2 | 0.336 |
| | | 3 | 0.336 |
| | 400 | 1 | 0.342 |
| | | 2 | 0.374 |
| | | 3 | 0.361 |
| | 1000 | 1 | 0.515 |
| | | 2 | 0.488 |
| | | 3 | 0.524 |
| **Population ≥ 10 trees (176 populations)** | 100 | 1 | 0.330 |
| | | 2 | 0.344 |
| | | 3 | 0.329 |
| | 400 | 1 | 0.325 |
| | | 2 | 0.340 |
| | | 3 | 0.334 |
| | **1000** | 1 | 0.527 |
| | | **2** | **0.538** |
| | | 3 | 0.529 |

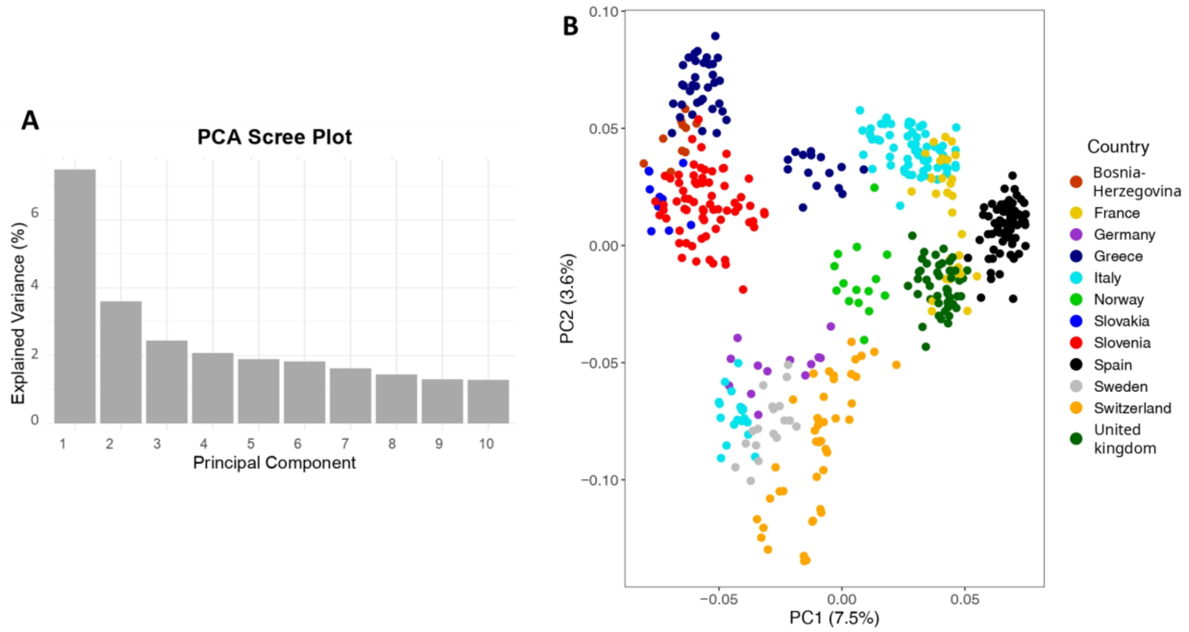


*Figure S4*. Principal component analysis (PCA) at the individual level using the SNP data. A: scree plot showing the percentage of explained genetic variance per principal component. B: Position of each individual on the first two axes of the PCA. Colours refer to the country of origin.

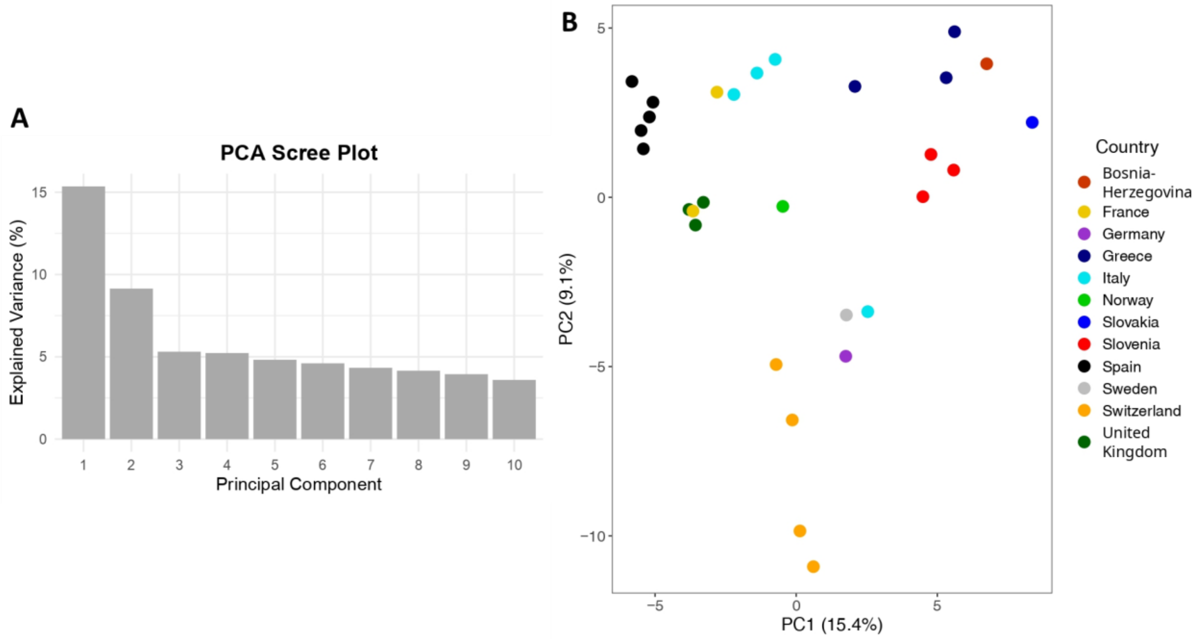


*Figure S5*. Principal component analysis (PCA) at the population level using SNP data. A: scree plot showing the percentage of explained genetic variance per principal component. B: Position of each population on the first two axes of the PCA. Colours refer to the country of origin.

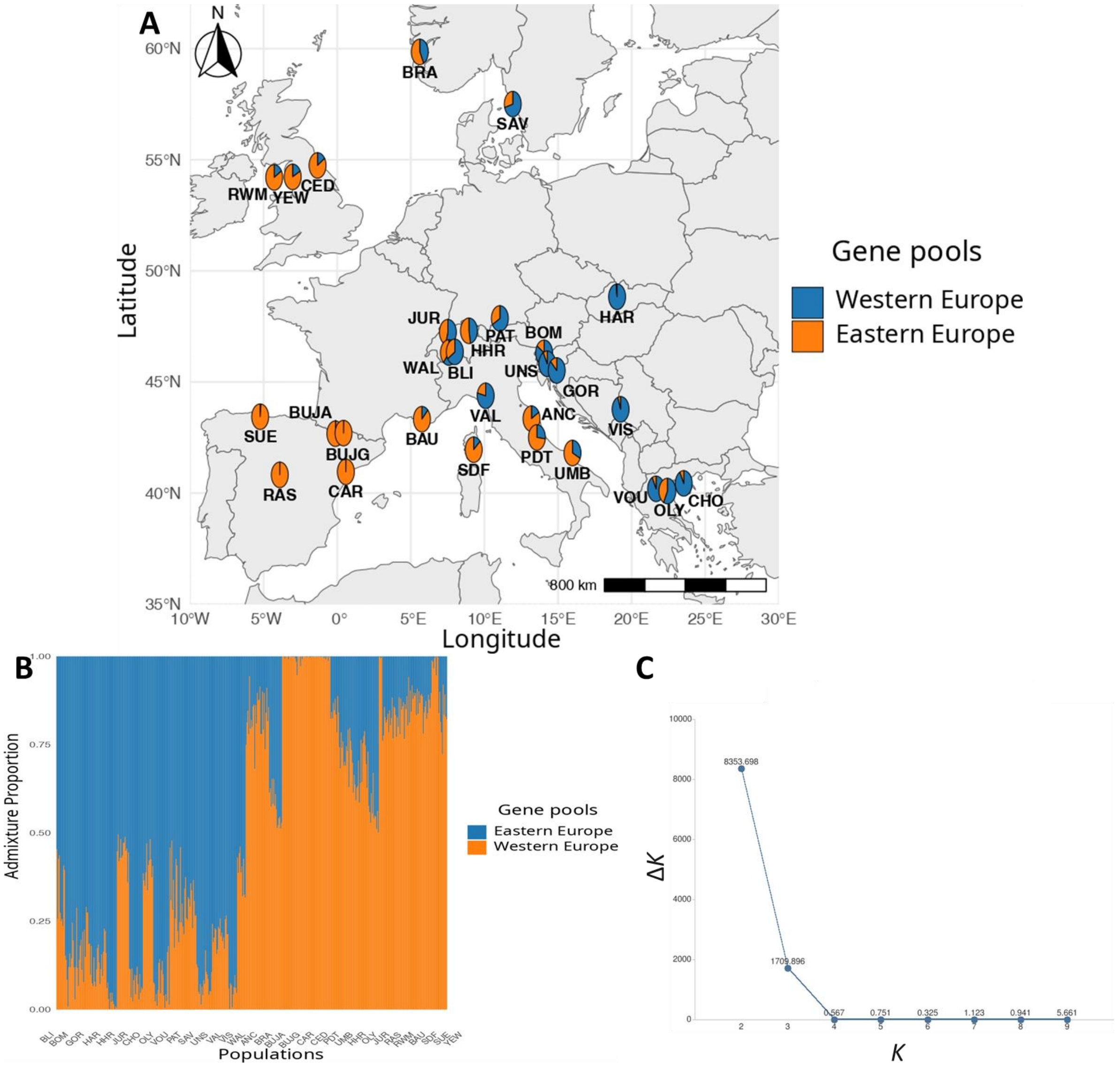


*Figure S6*. Population genetic structure analysis at the European geographical scale using the STRUCTURE Bayesian clustering approach. A: map showing ancestry coefficient pie charts for *K*=2, the best *K* (see C and main text). Each pie represents a population. B: bar chart showing ancestry coefficients for *K*=2. Each vertical line is an individual. C: Δ*K* results for *K* ranging from 2 to 9. Population labels correspond to those in Table S1.

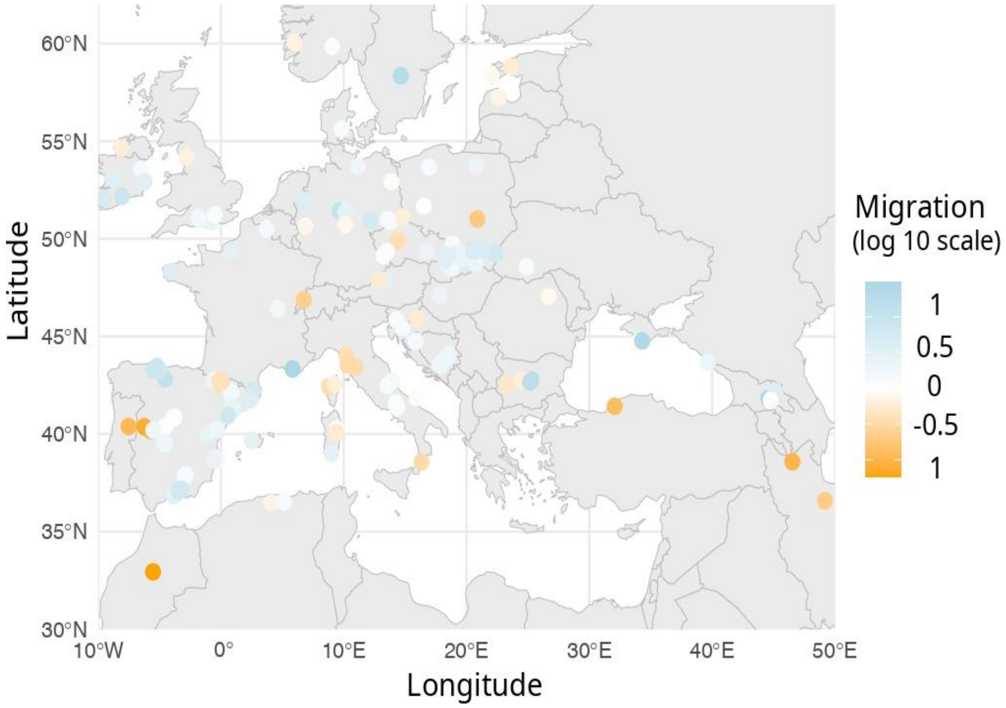


*Figure S7*. Historical effective migration at the European geographical scale obtained from the EEMS software. Positive values (in blue) indicate an estimated historical gene flow higher than expected under an isolation by distance (IBD) scenario, while negative values (in orange) indicate lower estimated historial gene flow than expected under IBD. The points represent the 176 nuSSR-genotyped populations retrieved from Mayol *et al*. (2015)[1] used to build the model.

[1]Mayol M, Riba M, González-Martínez SC, *et al.* (2015) Adapting through glacial cycles: insights from a long-lived tree (*Taxus baccata*). New Phytologist 208:973–986. https://doi.org/10.1111/nph.13496

### Section C: Patterns of local adaptation to climate

*Additional methods: details on GEA computations*

Several methods have been applied to *T. baccata* to compute gene-environment associations (GEA):

RDA and pRDA are linear multivariate analyses combining ordination and multiple regression approaches to produce canonical axes (Legendre and Legendre 2012). To account for population genetic structure in the pRDA, we used the first two axes of the PCA (see above and Figure S5b). Following Capblancq and Forester (2021), SNPs with an extreme Mahalanobis distance from the center of the RDA space along the significant axes were considered as candidate loci, with a false discovery rate (FDR) of 10% (Figure S8).

Latent Factor Mixed Model (LFMM) is a multivariate linear regression method that relies on least squares estimation to assess the effect size of each locus (Caye *et al*. 2019), and on latent factors to account for population genetic structure (two latent factors used in our study). An FDR of 10% was also applied here (Figure S9).

BayPass standard covariate (STD) model is a Bayesian univariate method that estimates the extent to which each marker is linearly associated with the predictors (Gautier 2015). Population genetic structure was accounted for using a variance-covariance matrix computed from the 8,252 SNPs not filtered by minor allele count (Figure S10). SNPs with mean Bayes factor (BF) > 8 over five independent runs, corresponding to “substantial evidence” on the Jeffreys’ scale (Kass and Raftery 1995), were considered candidate loci for climate adaptation (Figure S11).

Gradient Forest (GF) is a non-linear machine learning algorithm fitting an ensemble of regression trees to model changes in allele frequencies across populations (Ellis *et al*. 2012; Fitzpatrick *et al*. 2021). GF models were run on raw allele frequencies (GF_raw) or on allele frequencies corrected for population genetic structure (obtained by multiplying the matrix of effect sizes adjusted by LFMM by the transpose of the matrix of the climatic variables; GF_corrected). For each model, five independent runs were performed and for each run, the top 5% of the SNPs were retained; then, the overlapping top 5% of SNPs across the five independent runs were considered as candidate loci (Figure S12).

*References*


Capblancq T, Forester BR (2021) Redundancy analysis: A Swiss Army Knife for landscape genomics. Methods in Ecology and Evolution 12:2298–2309. https://doi.org/10.1111/2041-210X.13722

Caye K, Jumentier B, Lepeule J, François O (2019) LFMM 2: Fast and accurate inference of gene-environment associations in genome-wide studies. Molecular Biology and Evolution 36:852–860. https://doi.org/10.1093/molbev/msz008

Ellis N, Smith SJ, Pitcher CR (2012) Gradient forests: calculating importance gradients on physical predictors. Ecology 93:156–168. https://doi.org/10.1890/11-0252.1

Fitzpatrick MC, Chhatre VE, Soolanayakanahally RY, Keller SR (2021) Experimental support for genomic prediction of climate maladaptation using the machine learning approach Gradient Forests. Molecular Ecology Resources 21:2749–2765. https://doi.org/10.1111/1755-0998.13374

Gautier M (2015) Genome-wide scan for adaptive divergence and association with population-specific covariates. Genetics 201:1555–1579. https://doi.org/10.1534/genetics.115.181453

Legendre P, Legendre L (2012) Chapter 11 - Canonical analysis. In: Legendre P, Legendre L (eds) Developments in environmental modelling. Elsevier pp: 625–710.

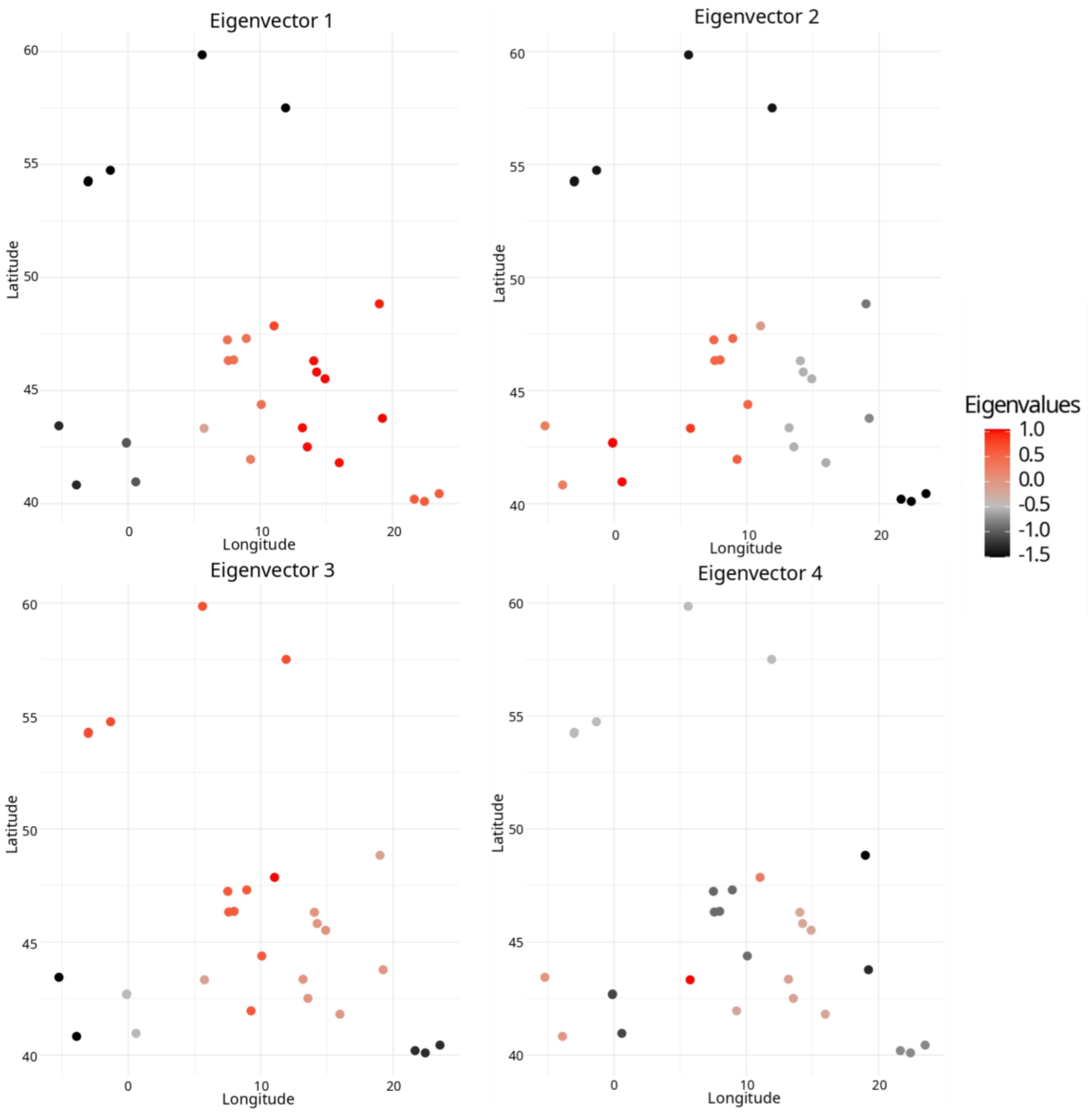


*Figure S8*. Calculation of distance-based Moran's eigenvectors (dbMEMs); the figure shows the maps for the four positive axes of spatial autocorrelation. Each point is a population and colour refers to the strength of the spatial autocorrelation, from -1.5 (negative autocorrelation in black) to one (positive autocorrelation in red).

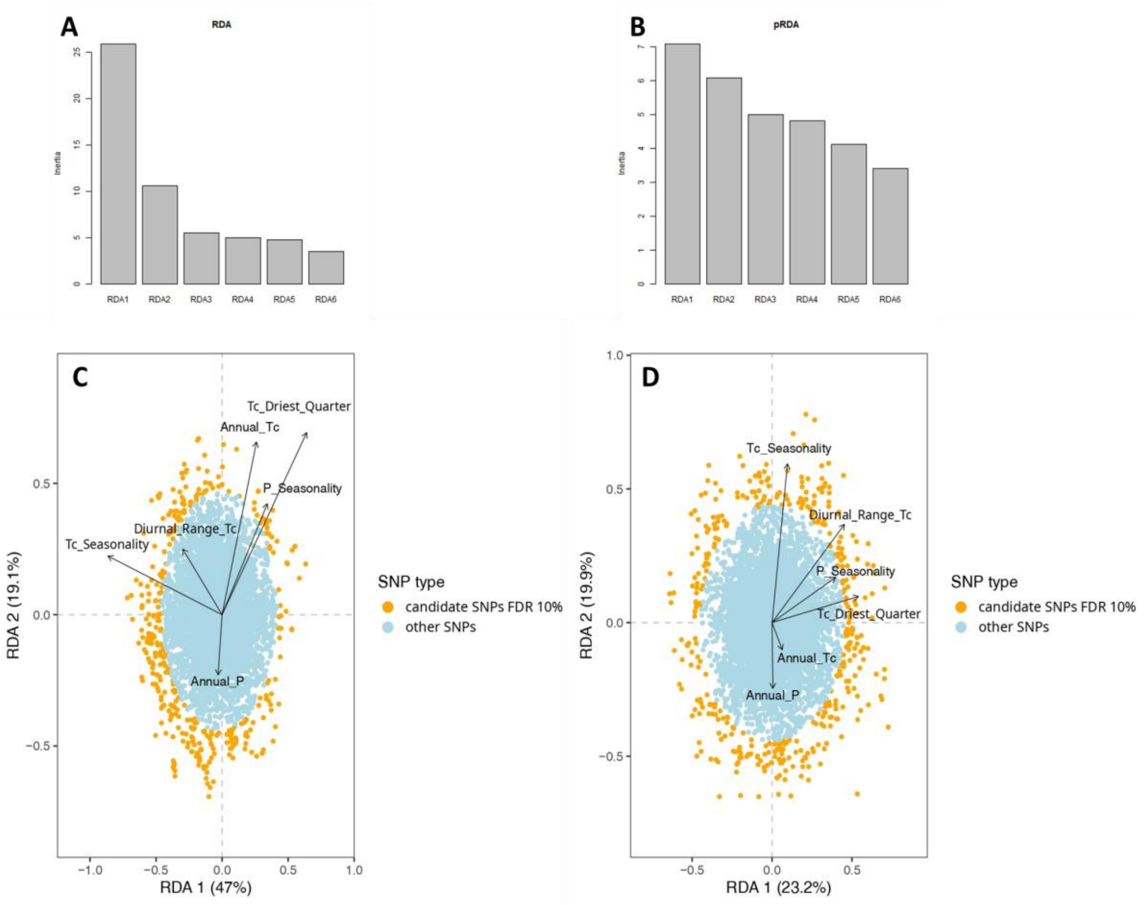


*Figure S9*. Detection of climate-associated loci using redundancy analysis (RDA) and partial redundancy analysis (pRDA) genotype-environment association (GEA) methods. A: scree plot displaying the inertia explained by each of the canonical axes for RDA and B: scree plot for pRDA. C and D: biplots showing the loci and the six climate predictors used in the GEA analyses for the two first canonical axes for the RDA (C) and the pRDA models (D). Each point is a SNP and colour refers to whether the SNP was identified or not  as associated with climatic predictors based on a false discovery rate (FDR) threshold of 10%.

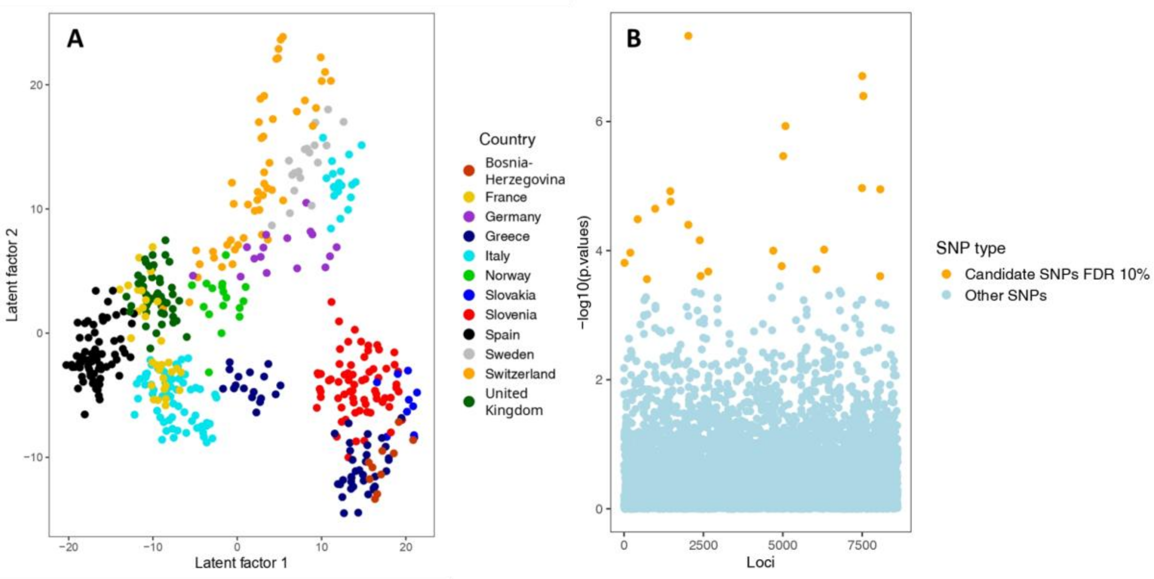


*Figure S10*. Detection of climate-associated loci using latent factor mixed model (LFMM) genotype-environment association (GEA) method. A: principal component analysis of the first two latent factors. Each point is an individual and colours refer to the country of origin. B: Manhattan plot displaying the loci based on their -log 10 *p*-value. Each point is a SNP and colour refers to whether the SNP was identified or not as associated with climatic predictors based on a false discovery rate (FDR) threshold of 10%.

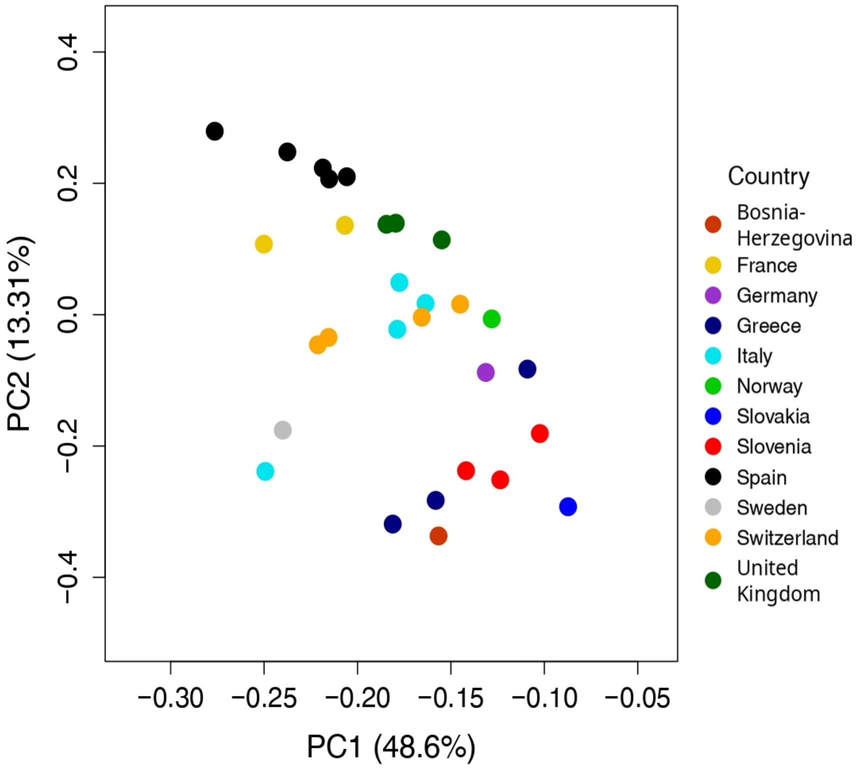


*Figure S11*. Principal component analysis (PCA) displaying the first two axes of the variance-covariance matrix (also referred to as omega matrix) obtained from the BayPass standard covariate (STD) model following Gautier (2015). Each point represents a population and colours refer to the country of origin.

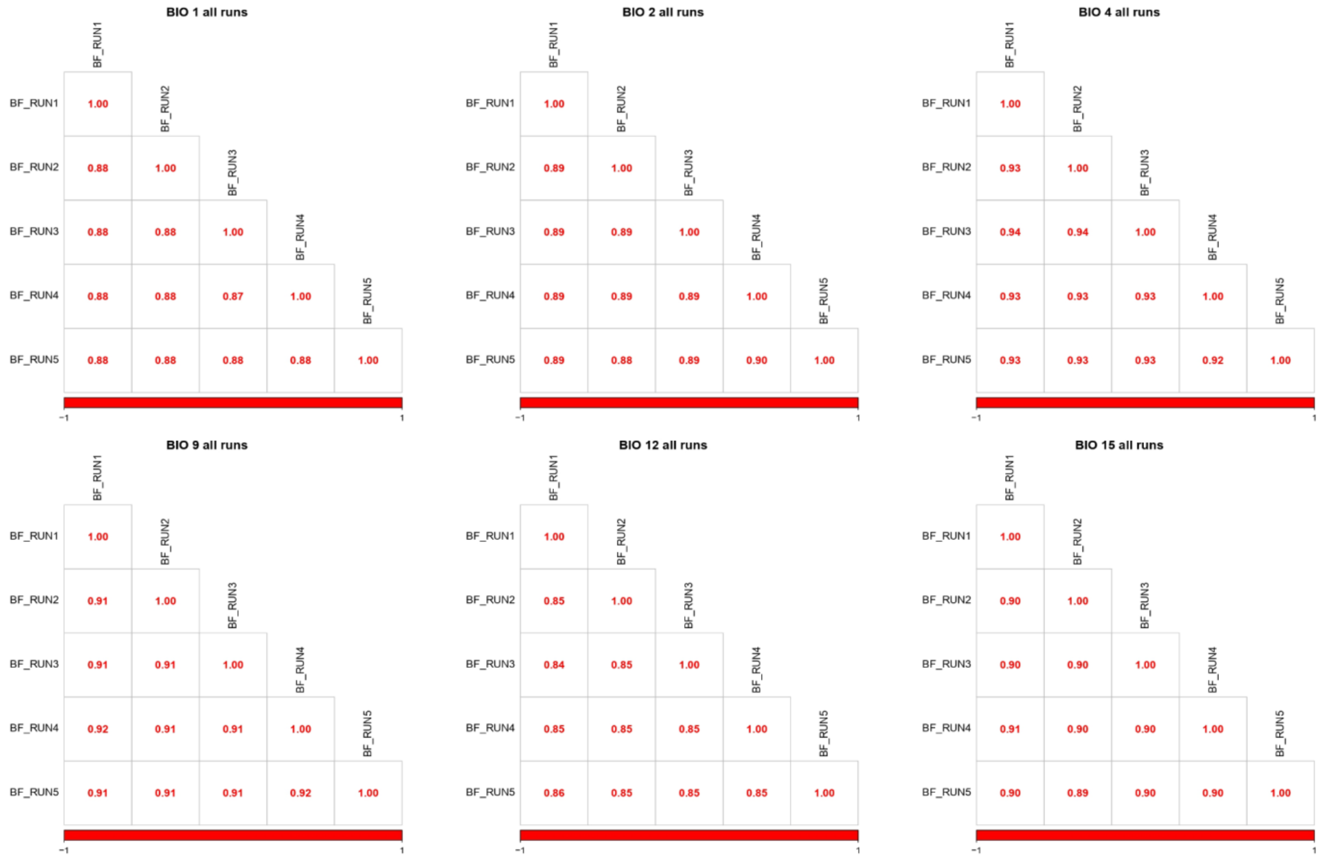


*Figure S12*. Detection of climate-associated loci using the BayPass genotype-environment association (GEA) method. Graphs show, for each of the six retained climate predictors, pairwise Pearson's correlations of the Bayes factor (BF) of each SNP across five independent runs of the BayPass model. Bio 1: Mean Annual Temperature; Bio 2: Mean Diurnal Range; Bio 4: Temperature Seasonality; Bio 9: Mean Temperature Driest Quarter; Bio 12: Mean Annual Precipitation and Bio 15: Precipitation Seasonality.

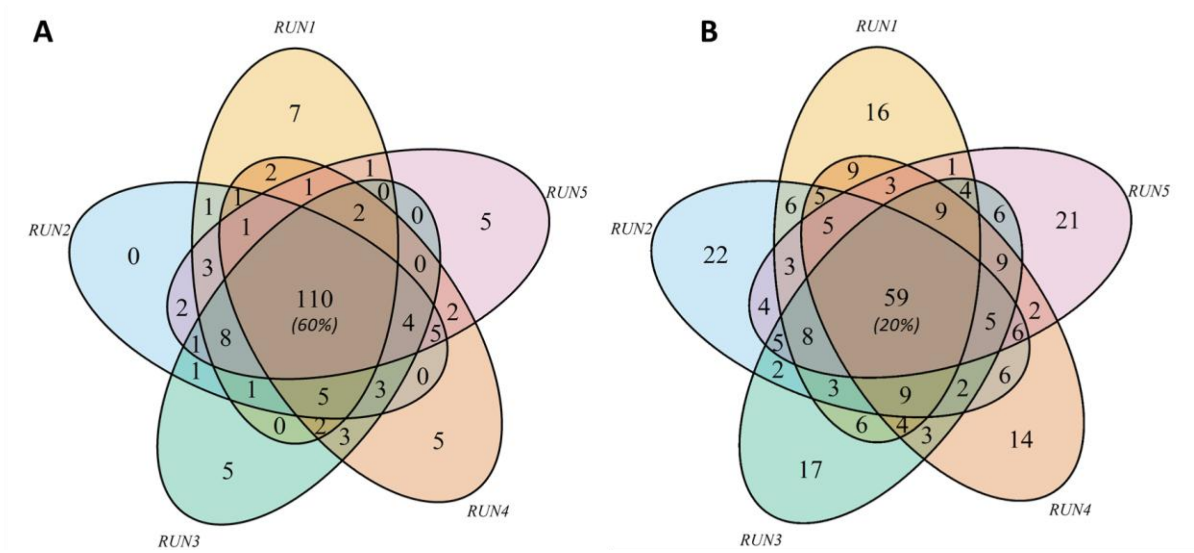


*Figure S13*. Venn diagrams showing the number of overlapping climate-associated loci across runs considering the top 5% of SNPs most associated with climatic predictors using the gradient forest genotype-environment association (GEA) method. A: model not corrected for population structure (GF-raw). B: model corrected for population structure (GF-corrected).

### Section D: Potential maladaptation under future climate

*Additional methods: details on Bayesian linear models*

For each model, we ran four MCMCs under a Gaussian distribution with 100,000 iterations each, including a burn-in period of 50,000 iterations, followed by a thinning interval of 50 iterations, using a regularising prior of N(0,1) for the coefficient of the predictor, selected after a prior predictive check analysis. The convergence of the chains was assessed using the Gelman-Rubin criteria (Gelman and Rubin 1992). To assess the goodness of fit of the model, the distribution of the observed response variable was visually compared with the simulated distribution obtained by the model using the pp_check function of the brms package (Bürkner 2021).

*References*

Bürkner P-C (2021) Bayesian item response modeling in R with brms and Stan. Journal of Statistical Software 100:1–54. https://doi.org/10.18637/jss.v100.i05

Gelman A, Rubin DB (1992) Inference from iterative simulation using multiple sequences. Statistical Science 7:457–472. https://doi.org/10.1214/ss/1177011136

*Table S8.* Population estimates for the genomic discrepancy index (GDI-RDA) and genomic offsets (computed using RDA and GF) for the 29 SNP-genotyped populations. All three indexes have been standardised using min-max normalization. Population labels correspond to those in Table S1.

| Population | Genomic discrepancy index (GDI-RDA) | Genomic offset (RDA) | Genomic offset (GF) |
|---|---|---|---|
| VIS | 0.59 | 0.44 | 0.43 |
| BAU | 0.11 | 1.00 | 0.14 |
| SDF | 0.49 | 0.63 | 0.34 |
| PAT | 0.10 | 0.23 | 0.30 |
| CHO | 0.99 | 0.55 | 0.49 |
| VOU | 0.32 | 0.64 | 0.60 |
| OLY | 1.00 | 0.75 | 0.56 |
| ANC | 0.69 | 0.60 | 0.58 |
| PDT | 0.03 | 0.36 | 0.72 |
| UMB | 0.60 | 0.33 | 0.26 |
| VAL | 0.76 | 0.60 | 0.42 |
| BRA | 0.04 | 0.20 | 0.05 |
| HAR | 0.43 | 0.00 | 0.20 |
| GOR | 0.14 | 0.19 | 0.38 |
| BOM | 0.12 | 0.56 | 0.42 |
| UNS | 0.24 | 0.47 | 0.48 |
| BUJA | 0.47 | 0.70 | 0.63 |
| BUJG | 0.43 | 0.70 | 0.65 |
| CAR | 0.52 | 0.68 | 0.49 |
| RAS | 0.24 | 0.43 | 1.00 |
| SUE | 0.03 | 0.01 | 0.04 |
| SAV | 0.00 | 0.65 | 0.34 |
| BLI | 0.56 | 0.67 | 0.47 |
| HHR | 0.11 | 0.40 | 0.32 |
| JUR | 0.23 | 0.20 | 0.30 |
| WAL | 0.58 | 0.51 | 0.47 |
| CED | 0.34 | 0.19 | 0.01 |
| RWM | 0.22 | 0.06 | 0.00 |
| YEW | 0.05 | 0.05 | 0.00 |

*Table S9*. Summary of Pearson's correlations (*r*) assessing the association between genomic offset models and fitness proxies. *P*-values were calculated using Student's t-test. Significant associations based on *p*-values < 0.05 are highlighted in bold.

| Method | Model | Trait | *r* | *p*-value |
|---|---|---|---|---|
| RDA | All | Growth | -0.302 | 0.152 |
| RDA | All | Growth phenology | -0.108 | 0.632 |
| RDA | All | Reproductive phenology | -0.046 | 0.876 |
| RDA | All | Leaf thickness | -0.225 | 0.327 |
| RDA | All | Composite index | -0.317 | 0.131 |
| RDA | Random | Growth | -0.335 | 0.110 |
| RDA | Random | Growth phenology | -0.213 | 0.341 |
| RDA | Random | Reproductive phenology | -0.096 | 0.745 |
| RDA | Random | Leaf thickness | -0.351 | 0.118 |
| **RDA** | **Random** | **Composite index** | **-0.439** | **0.032** |
| RDA | Random_2 | Growth | -0.273 | 0.197 |
| RDA | Random_2 | Growth phenology | -0.123 | 0.585 |
| RDA | Random_2 | Reproductive phenology | -0.020 | 0.946 |
| RDA | Random_2 | Leaf thickness | -0.186 | 0.420 |
| RDA | Random_2 | Composite index | -0.298 | 0.157 |
| RDA | Outlier | Growth | -0.297 | 0.159 |
| RDA | Outlier | Growth phenology | -0.246 | 0.270 |
| RDA | Outlier | Reproductive phenology | -0.193 | 0.510 |
| RDA | Outlier | Leaf thickness | -0.282 | 0.215 |
| **RDA** | **Outlier** | **Composite index** | **-0.421** | **0.041** |
| RDA | all_outlier | Growth | -0.314 | 0.135 |
| RDA | all_outlier | Growth phenology | -0.124 | 0.582 |
| RDA | all_outlier | Reproductive phenology | -0.052 | 0.860 |
| RDA | all_outlier | Leaf thickness | -0.255 | 0.265 |
| RDA | all_outlier | Composite index | -0.333 | 0.112 |
| GF | All | Growth | -0.277 | 0.191 |
| GF | All | Growth phenology | -0.288 | 0.193 |
| GF | All | Reproductive phenology | -0.333 | 0.244 |
| GF | All | Leaf thickness | -0.147 | 0.525 |
| GF | All | Composite index | -0.395 | 0.056 |
| GF | Random | Growth | -0.304 | 0.148 |
| GF | Random | Growth phenology | -0.325 | 0.140 |
| GF | Random | Reproductive phenology | -0.348 | 0.223 |
| GF | Random | Leaf thickness | -0.228 | 0.320 |
| **GF** | **Random** | **Composite index** | **-0.495** | **0.014** |
| GF | Random_2 | Growth | -0.309 | 0.141 |
| GF | Random_2 | Growth phenology | -0.299 | 0.175 |
| GF | Random_2 | Reproductive phenology | -0.312 | 0.278 |
| GF | Random_2 | Leaf thickness | -0.169 | 0.465 |
| **GF** | **Random_2** | **Composite index** | **-0.421** | **0.040** |
| GF | Outlier | Growth | -0.279 | 0.187 |
| GF | Outlier | Growth phenology | -0.260 | 0.243 |

| | | | | |
|---|---|---|---|---|
| GF | Outlier | Reproductive phenology | -0.302 | 0.293 |
| GF | Outlier | Leaf thickness | -0.258 | 0.258 |
| GF | Outlier | Composite index | -0.402 | 0.052 |
| GF | all_outlier | Growth | -0.286 | 0.175 |
| GF | all_outlier | Growth phenology | -0.293 | 0.186 |
| GF | all_outlier | Reproductive phenology | -0.318 | 0.268 |
| GF | all_outlier | Leaf thickness | -0.213 | 0.354 |
| **GF** | **all_outlier** | **Composite index** | **-0.421** | **0.040** |

*Table S10*. Summary of the frequentist linear models assessing the association between the genomic offset models and the fitness proxies. Significant associations based on *p*-values < 0.05 are highlighted in bold.

| Method | Model | Trait | $R^2$ | Intercept | Coeff. | *p*-value |
|---|---|---|---|---|---|---|
| RDA | All | Growth | 0.062 | 0.103 | -2.184 | 0.220 |
| RDA | All | Growth phenology | 0.005 | 0.003 | -0.072 | 0.726 |
| RDA | All | Reproductive phenology | 0.000 | 0.003 | -0.005 | 0.986 |
| RDA | All | Leaf thickness | 0.071 | 0.009 | -0.104 | 0.209 |
| RDA | All | Composite index | 0.092 | 0.546 | -1.129 | 0.149 |
| RDA | Random | Growth | 0.038 | 0.077 | -2.269 | 0.339 |
| RDA | Random | Growth phenology | 0.001 | 0.000 | -0.050 | 0.853 |
| RDA | Random | Reproductive phenology | 0.001 | 0.000 | 0.048 | 0.903 |
| RDA | Random | Leaf thickness | 0.140 | 0.013 | -0.198 | 0.071 |
| RDA | Random | Composite index | 0.074 | 0.547 | -1.386 | 0.198 |
| RDA | Random_2 | Growth | 0.075 | 0.087 | -2.437 | 0.197 |
| RDA | Random_2 | Growth phenology | 0.015 | 0.005 | -0.134 | 0.585 |
| RDA | Random_2 | Reproductive phenology | 0.000 | 0.005 | -0.026 | 0.946 |
| RDA | Random_2 | Leaf thickness | 0.030 | 0.006 | -0.076 | 0.452 |
| RDA | Random_2 | Composite index | 0.072 | 0.544 | -1.147 | 0.206 |
| RDA | Outlier | Growth | 0.071 | 0.093 | -2.149 | 0.188 |
| RDA | Outlier | Growth phenology | 0.031 | 0.009 | -0.160 | 0.389 |
| RDA | Outlier | Reproductive phenology | 0.009 | 0.009 | -0.109 | 0.707 |
| RDA | Outlier | Leaf thickness | 0.143 | 0.011 | -0.133 | 0.069 |
| **RDA** | **Outlier** | **Composite index** | **0.189** | **0.568** | **-1.478** | **0.034** |
| RDA | all_outlier | Growth | 0.098 | 0.088 | -2.370 | 0.135 |
| RDA | all_outlier | Growth phenology | 0.015 | 0.004 | -0.116 | 0.582 |
| RDA | all_outlier | Reproductive phenology | 0.003 | 0.007 | -0.064 | 0.860 |
| RDA | all_outlier | Leaf thickness | 0.058 | 0.008 | -0.085 | 0.294 |
| RDA | all_outlier | Composite index | 0.101 | 0.547 | -1.095 | 0.131 |
| GF | All | Growth | 0.062 | 0.084 | -1.671 | 0.219 |
| GF | All | Growth phenology | 0.047 | 0.011 | -0.163 | 0.289 |
| GF | All | Reproductive phenology | 0.025 | 0.014 | -0.152 | 0.532 |
| GF | All | Leaf thickness | 0.098 | 0.009 | -0.090 | 0.136 |
| **GF** | **All** | **Composite index** | **0.175** | **0.560** | **-1.160** | **0.042** |
| GF | Random | Growth | 0.071 | 0.156 | -13.606 | 0.187 |
| GF | Random | Growth phenology | 0.069 | 0.021 | -1.515 | 0.194 |
| GF | Random | Reproductive phenology | 0.062 | 0.029 | -1.730 | 0.317 |
| GF | Random | Leaf thickness | 0.036 | 0.008 | -0.417 | 0.376 |
| GF | Random | Composite index | 0.153 | 0.595 | -8.303 | 0.059 |
| GF | Random_2 | Growth | 0.096 | 0.150 | -14.895 | 0.141 |
| GF | Random_2 | Growth phenology | 0.089 | 0.026 | -1.827 | 0.176 |
| GF | Random_2 | Reproductive phenology | 0.097 | 0.038 | -2.237 | 0.278 |
| GF | Random_2 | Leaf thickness | 0.029 | 0.007 | -0.373 | 0.460 |
| **GF** | **Random_2** | **Composite index** | **0.179** | **0.612** | **-9.174** | **0.039** |
| GF | Outlier | Growth | 0.081 | 0.163 | -10.958 | 0.159 |
| GF | Outlier | Growth phenology | 0.064 | 0.020 | -1.099 | 0.213 |

| | | | | | | |
|---|---|---|---|---|---|---|
| GF | Outlier | Reproductive phenology | 0.092 | 0.036 | -1.784 | 0.221 |
| GF | Outlier | Leaf thickness | 0.069 | 0.010 | -0.435 | 0.215 |
| **GF** | **Outlier** | **Composite index** | **0.195** | **0.606** | **-7.056** | **0.031** |
| GF | all_outlier | Growth | 0.082 | 0.087 | -7.173 | 0.175 |
| GF | all_outlier | Growth phenology | 0.086 | 0.019 | -0.914 | 0.186 |
| GF | all_outlier | Reproductive phenology | 0.101 | 0.034 | -1.297 | 0.268 |
| GF | all_outlier | Leaf thickness | 0.044 | 0.007 | -0.238 | 0.361 |
| **GF** | **all_outlier** | **Composite index** | **0.175** | **0.582** | **-4.751** | **0.042** |

*Table S11*. Summary of the Bayesian linear models assessing the association between the genomic offset models and the fitness proxies. Rhat: R-hat convergence diagnostic; CI: credible intervals; *b:* mean value of the posterior distribution for the linear coefficient. Credible intervals of coefficient values that do not include zero are highlighted in bold.

| **Method** | **Model** | **Trait** | ***Rhat*** | ***$R^2$*** | ***$R^2$* CI** | ***Coefficient*** | ***Coefficient CI*** |
|---|---|---|---|---|---|---|---|
| RDA | All | Growth | 1.000 | 0.080 | 0-0.266 | -0.237 | -0.626-0.155 |
| RDA | All | Growth phenology | 1.000 | 0.040 | 0-0.167 | -0.070 | -0.471-0.348 |
| RDA | All | Reproductive phenology | 1.000 | 0.050 | 0-0.208 | -0.001 | -0.521-0.496 |
| RDA | All | Leaf thickness | 1.001 | 0.090 | 0-0.292 | -0.253 | -0.672-0.174 |
| RDA | All | Composite index | 1.001 | 0.104 | 0-0.325 | -0.285 | -0.723-0.135 |
| RDA | Random | Growth | 1.001 | 0.089 | 0-0.286 | -0.253 | -0.659-0.164 |
| RDA | Random | Growth phenology | 1.000 | 0.060 | 0-0.235 | -0.174 | -0.593-0.235 |
| RDA | Random | Reproductive phenology | 1.000 | 0.056 | 0-0.243 | -0.089 | -0.636-0.426 |
| RDA | Random | Leaf thickness | 1.000 | 0.146 | 0.001-0.363 | -0.363 | -0.755-0.035 |
| **RDA** | **Random** | **Composite index** | **1.000** | **0.183** | **0.004-0.412** | **-0.418** | **-0.821--0.037** |
| RDA | Random_2 | Growth | 1.001 | 0.092 | 0-0.297 | -0.259 | -0.678-0.161 |
| RDA | Random_2 | Growth phenology | 1.000 | 0.054 | 0-0.221 | -0.120 | -0.574-0.352 |
| RDA | Random_2 | Reproductive phenology | 1.000 | 0.063 | 0-0.249 | -0.016 | -0.610-0.600 |
| RDA | Random_2 | Leaf thickness | 1.000 | 0.065 | 0-0.248 | -0.166 | -0.622-0.295 |
| RDA | Random_2 | Composite index | 1.000 | 0.091 | 0-0.294 | -0.255 | -0.675-0.168 |
| RDA | Outlier | Growth | 1.001 | 0.081 | 0-0.285 | -0.233 | -0.664-0.194 |
| RDA | Outlier | Growth phenology | 1.000 | 0.070 | 0-0.252 | -0.207 | -0.612-0.194 |
| RDA | Outlier | Reproductive phenology | 1.000 | 0.068 | 0-0.267 | -0.152 | -0.671-0.376 |
| RDA | Outlier | Leaf thickness | 1.000 | 0.113 | 0-0.331 | -0.306 | -0.729-0.099 |
| **RDA** | **Outlier** | **Composite index** | **1.000** | **0.172** | **0.002-0.398** | **-0.403** | **-0.801--0.004** |
| RDA | all_outlier | Growth | 1.000 | 0.114 | 0-0.331 | 305 | -0.728-0.126 |
| RDA | all_outlier | Growth phenology | 1.001 | 0.052 | 0-0.212 | -0.121 | -0.558-0.340 |

| | | | | | | | |
|---|---|---|---|---|---|---|---|
| RDA | all_outlier | Reproductive phenology | 1.000 | 0.064 | 0-0.249 | -0.047 | -0.644-0.554 |
| RDA | all_outlier | Leaf thickness | 1.001 | 0.082 | 0-0.283 | -0-224 | -0.666-0.233 |
| RDA | all_outlier | Composite index | 1.000 | 0.112 | 0-0.321 | -0.302 | -0.705-0.108 |
| GF | All | Growth | 1.000 | 0.088 | 0-0.284 | -0.255 | -0.654-0.136 |
| GF | All | Growth phenology | 1.000 | 0.089 | 0-0.288 | -0.256 | -0.664-0.158 |
| GF | All | Reproductive phenology | 1.000 | 0.091 | 0-0.320 | -0.235 | -0.746-0.270 |
| GF | All | Leaf thickness | 1.000 | 0.064 | 0-0.250 | -0.177 | -0.618-0.255 |
| GF | All | Composite index | 0.999 | 0.152 | 0.001-0.375 | -0.372 | -0.771-0.023 |
| GF | Random | Growth | 1.000 | 0.078 | 0-0.266 | -0.231 | -0.630-0.166 |
| GF | Random | Growth phenology | 1.000 | 0.106 | 0-0.304 | -0.293 | -0.675-0.099 |
| GF | Random | Reproductive phenology | 1.000 | 0.120 | 0-0.356 | -0.304 | -0.788-0.184 |
| GF | Random | Leaf thickness | 1.000 | 0.082 | 0-0.274 | -0.236 | -0.645-0.189 |
| **GF** | **Random** | **Composite index** | **1.000** | **0.232** | **0.012-0.456** | **-0.479** | **-0.862--0.091** |
| GF | Random_2 | Growth | 1.002 | 0.107 | 0-0.326 | -0.292 | -0.719-0.121 |
| GF | Random_2 | Growth phenology | 1.000 | 0.105 | 0-0.318 | -0.279 | -0.708-0.165 |
| GF | Random_2 | Reproductive phenology | 1.001 | 0.123 | 0-0.391 | -0.297 | -0.891-0.276 |
| GF | Random_2 | Leaf thickness | 1.000 | 0.063 | 0-0.243 | -0.160 | -0.609-0.318 |
| GF | Random_2 | Composite index | 1.000 | 0.175 | 0.003-0.404 | -0.407 | -0.812-0.002 |
| GF | Outlier | Growth | 1.000 | 0.092 | 0-0.286 | -0.262 | -0.653-0.147 |
| GF | Outlier | Growth phenology | 1.000 | 0.082 | 0-0.277 | -0.238 | -0.647-0.173 |
| GF | Outlier | Reproductive phenology | 1.000 | 0.088 | 0-0.314 | -0.223 | -0.739-0.294 |
| GF | Outlier | Leaf thickness | 0.999 | 0.095 | 0-0.298 | -0.267 | -0.679-0.153 |
| GF | Outlier | Composite index | 1.001 | 0.158 | 0.002-0.385 | -0.379 | -0.787-0.028 |
| GF | all_outlier | Growth | 1.001 | 0.096 | 0-0.308 | -0.265 | -0.695-0.164 |
| GF | all_outlier | Growth phenology | 1.000 | 0.100 | 0-0.310 | -0.271 | -0.696-0.169 |
| GF | all_outlier | Reproductive phenology | 1.001 | 0.122 | 0-0.388 | -0.288 | -0.875-0.278 |
| GF | all_outlier | Leaf thickness | 1.000 | 0.073 | 0-0.280 | -0.194 | -0.675-0.274 |
| GF | all_outlier | Composite index | 1.001 | 0.172 | 0.002-0.397 | -0.403 | -0.800-0.003 |

*Table S12*. Top BLAST hits for candidate genes with ≥ 3 SNPs per gene. Candidate genes were annotated using BLASTX searches against the NCBI non-redundant protein sequence database. In bold, the two genes that were identified as under selection in Mayol *et al*. (2020)[1].

| Gene code | Top BLAST hit | Function |
|---|---|---|
| **Tbac_129399** | Early-responsive to dehydration stress protein | Response to drought stress |
| **Tbac_72122** | Microtubule-severing ATPase | Microtubule cytoskeleton organisation |
| Tbac_19521 | Subtilisin-like protease SBT1.3 | Serine protease; proteolysis |
| Tbac_19904 | Calcium-dependent protein kinase | Intracellular signal transduction |
| Tbac_22150 | PPM-type phosphatase domain-containing protein | Developmental protein |

[1]Mayol M, Riba M, Cavers S, *et al.* (2020) A multiscale approach to detect selection in nonmodel tree species: Widespread adaptation despite population decline in *Taxus baccata* L. Evolutionary Applications 13:143–160. https://doi.org/10.1111/eva.12838

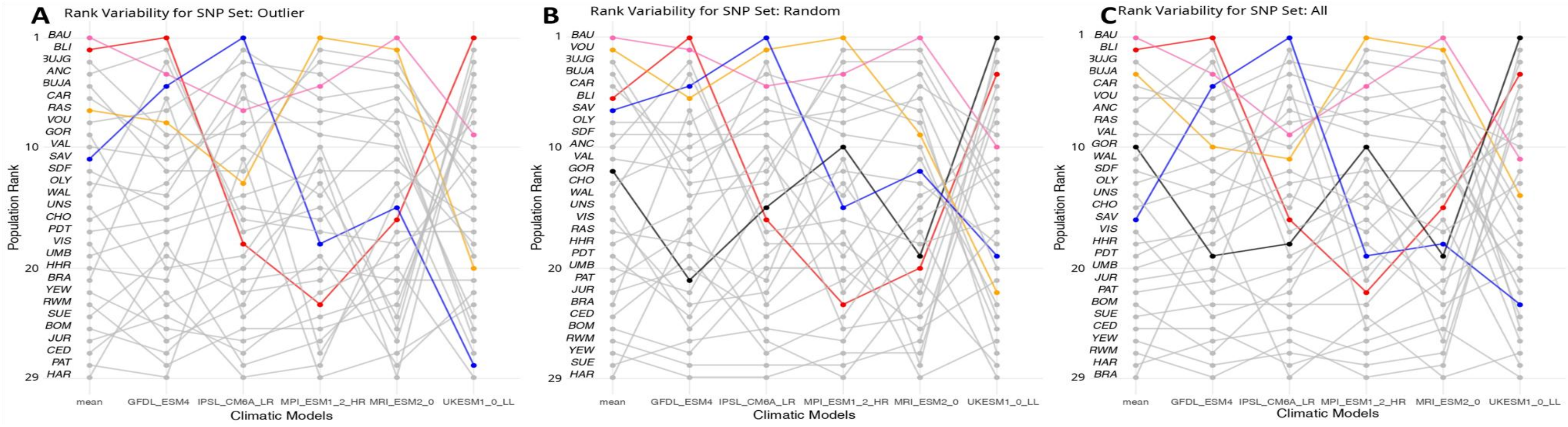


*Figure S14*. Population rank based on genomic offset predictions with RDA models using three SNP sets (*outlier*: 100 climate-associated loci, *random*: 100 random markers matching the allelic frequency of the outlier SNPs, and *all*: all the 8,616 SNPs) under the socio-economic pathway SSP 3-7.0 for the 2041-2070 time interval and five global climate models (GCMs). Mean predictions across the five GCMs are also provided. Ranks range from one (highest genomic offset) to 29 (lowest genomic offset). Coloured populations correspond to the populations having a rank of one in any of the models. Population order corresponds to the rank based on the mean genomic offset predictions. A: population rank for the *outlier* set; B: *random* set; and C: *all* loci.

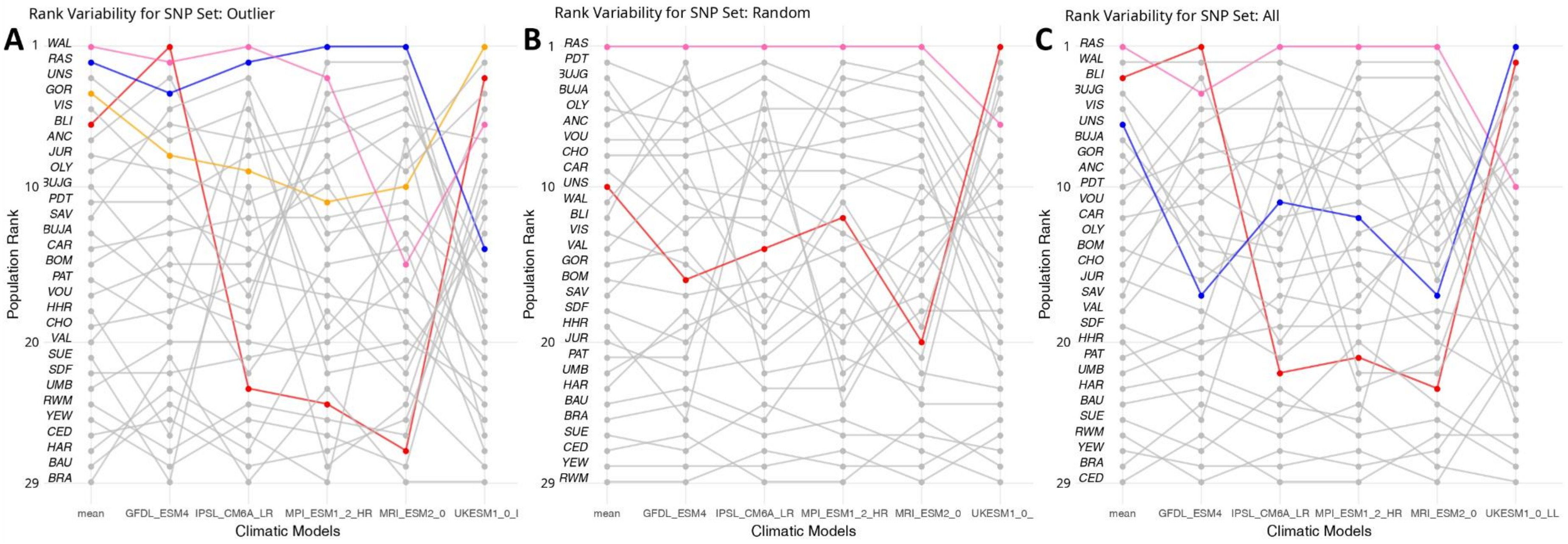


*Figure S15*. Population rank based on genomic offset predictions with GF models using three SNP sets (*outlier*: 100 climate-associated loci, *random*: 100 random markers matching the allelic frequency of the outlier SNPs, and *all*: all the 8,616 SNPs) under the socio-economic pathway SSP 3-7.0 for the 2041-2070 time interval and five global climate models (GCMs). Mean predictions across the five GCMs are also provided. Ranks range from one (highest genomic offset) to 29 (lowest genomic offset). Coloured populations correspond to the populations having a rank of one in any model. Population order corresponds to the rank based on the mean genomic offset predictions. A: population rank for the *outlier* set; B: *random* set; and C: *all* loci.

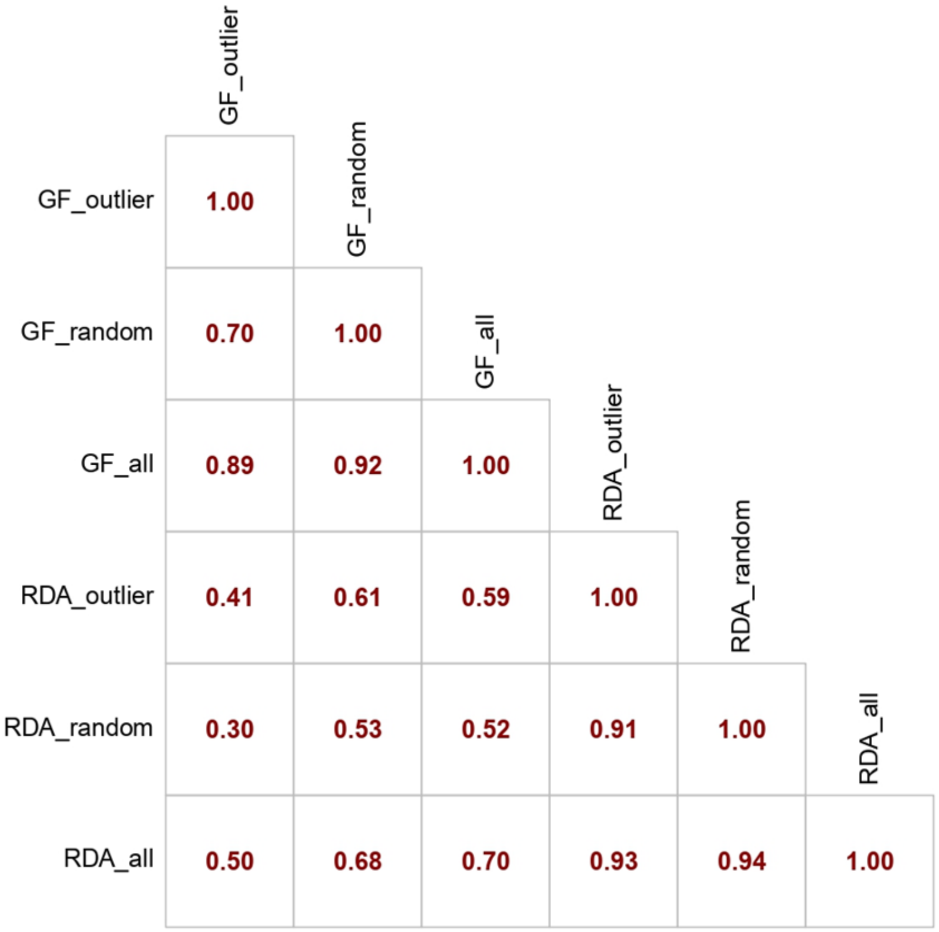


*Figure S16*. Pearson's correlation matrix of the mean genomic offset predictions across the five global climate models (GCMs) for GF and RDA models using three SNP sets (*outlier*: 100 climate-associated loci, *random*: 100 random markers matching the allelic frequency of the outlier SNPs, and *all*: all the 8,616 SNPs) under the socio-economic pathway SSP 3-7.0 for the 2041-2070 time interval.

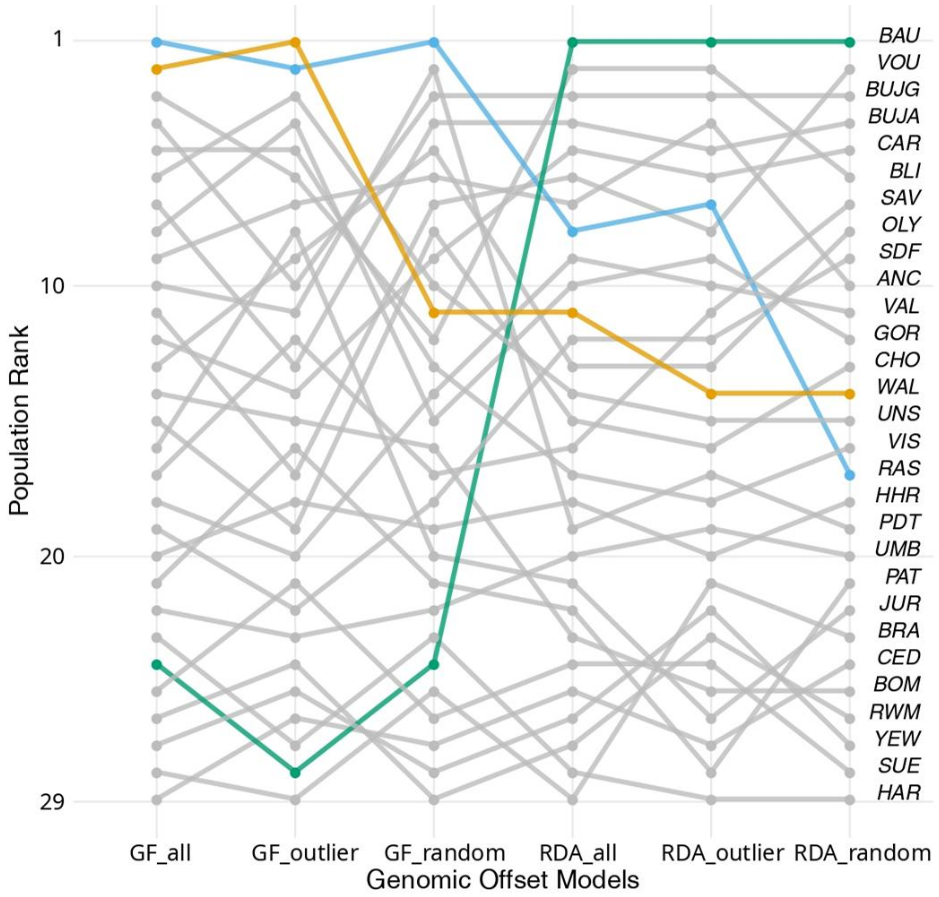


*Figure S17*. Population rank based on genomic offset predictions with GF and RDA models using three SNP sets (*outlier*: 100 climate-associated loci, *random*: 100 random markers matching the allelic frequency of the outlier SNPs, and *all*: all the 8,616 SNPs), under the socio-economic pathway SSP 3-7.0 for the 2041-2070 time interval. Ranks correspond to the mean predictions across five global climate models (GCMs) and range from one (highest genomic offset) to 29 (lowest genomic offset). Coloured populations correspond to the populations having a rank of one in any of the six models. Population order corresponds to the rank based on the RDA model with the *random* set of loci.

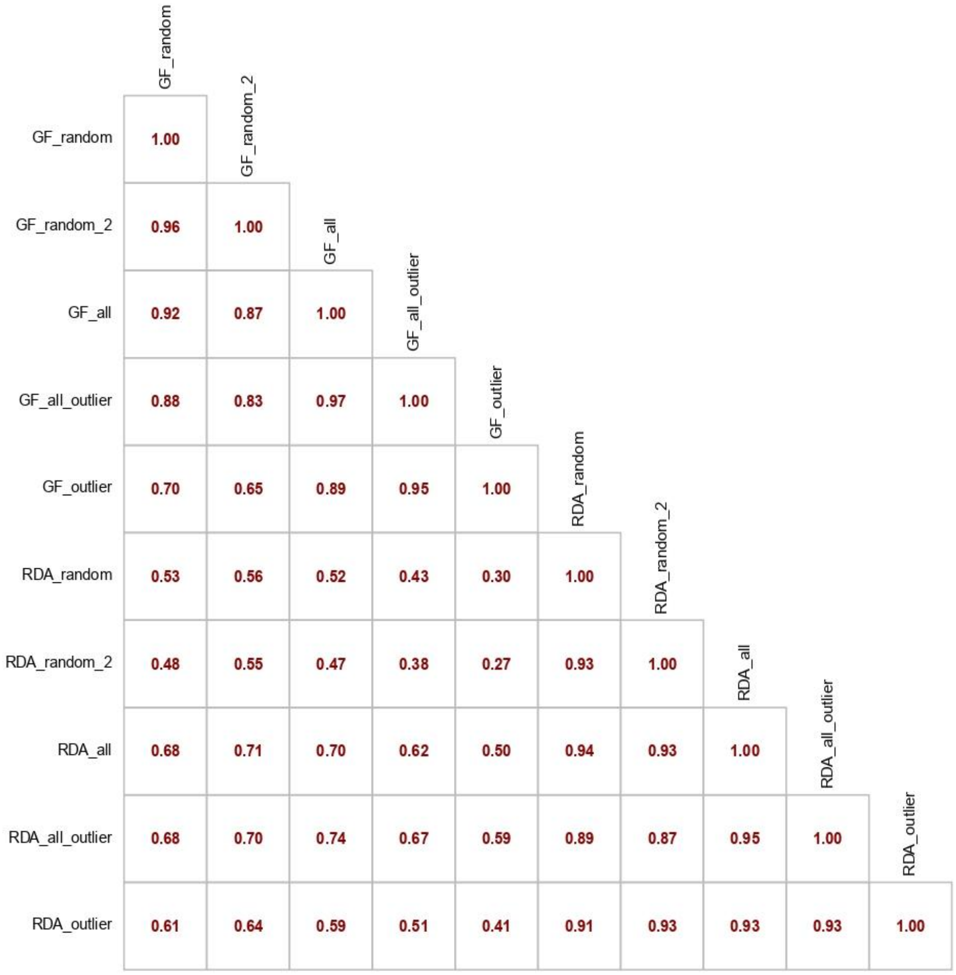


*Figure S18*. Pearson's correlation matrix of the mean genomic offset predictions across the five global climate models (GCMs) for GF and RDA models using all the SNP sets: *all*: all the 8,616 SNPs, *all_outlier*: 935 climate-associated loci identified by at least one GEA method, *outlier*: 100 climate-associated loci identified by at least two GEA methods, *random*: 100 random SNPs matching the allelic frequency of the outlier set, *random_2*: new set of 100 random SNPs matching the allelic frequency of the outlier set (only 4 SNPs overlapping with the *random* set), under the socio-economic pathway SSP 3-7.0 for the 2041-2070 time interval.

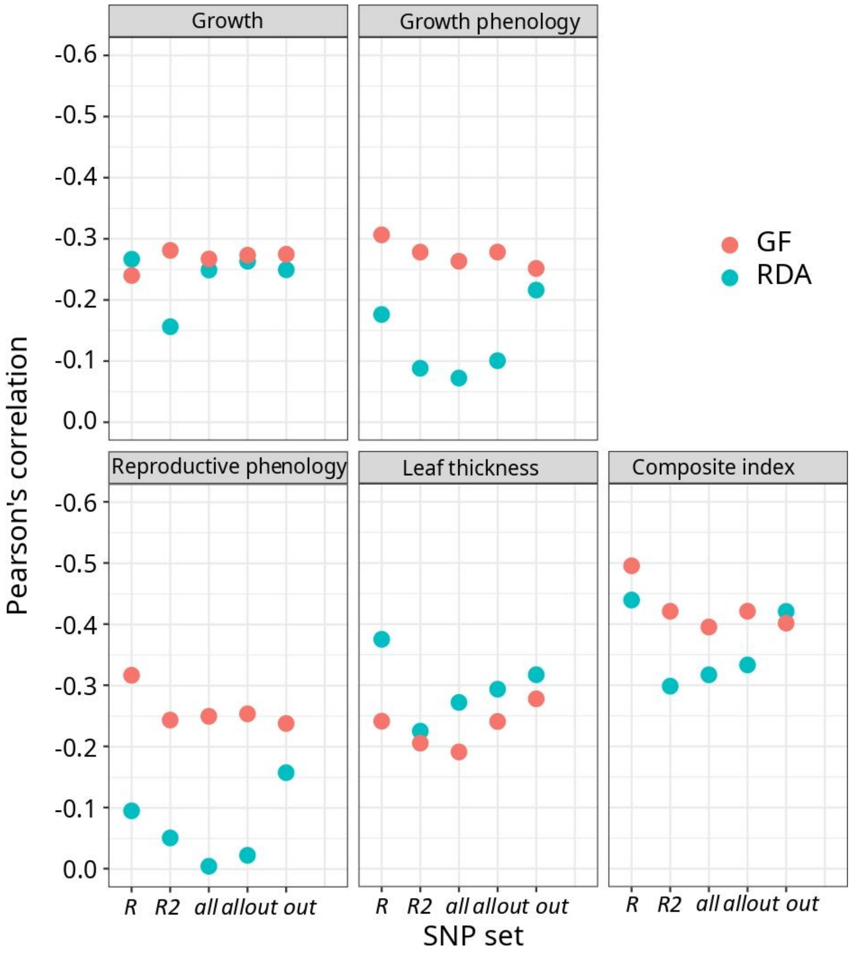


*Figure S19*. Pearson's correlations between the genomic offset models and the fitness components assessed in a comparative experiment under common garden conditions. Genomic offset was calculated for each population as the Euclidean distance between the predicted optimal genomic composition in the climate of origin of the population and that of the comparative experiment. The SNP sets are: *R* = *random*: 100 random SNPs matching the allelic frequency of the *outlier* set, *R2* = *random_2*: new set of 100 random SNPs matching the allelic frequency of the outlier set (only 4 SNPs overlapping with the *random* set), *all*: all the 8,616 SNPs, *allout* = *all_outlier*: 935 climate-associated loci identified by at least one GEA method, and *out* = *outlier*: 100 climate-associated loci identified by at least two GEA methods.

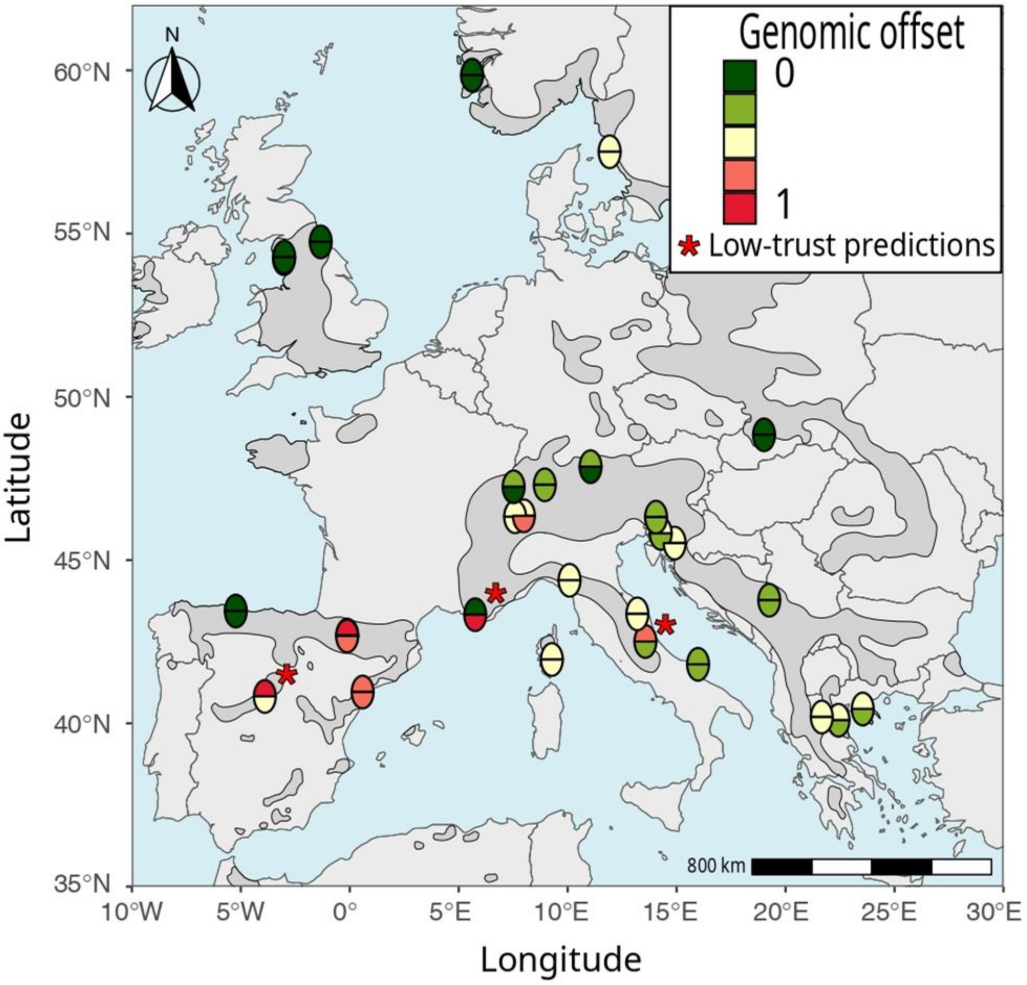


*Figure S20*. Genomic offset predictions for the RDA (lower half-circle) and GF (upper half-circle) models using the *random_2* set of SNPs for the mean climate across five GCMs for the 2041-2070 time interval under the SSP 3-7.0 using the 1901-1950 climate as reference period. Genomic offset predictions were standardised to range from zero (low - green) to one (high - red) and categorised into five classes. Red stars represent the populations where the predictions of the two models differ by more than two classes.

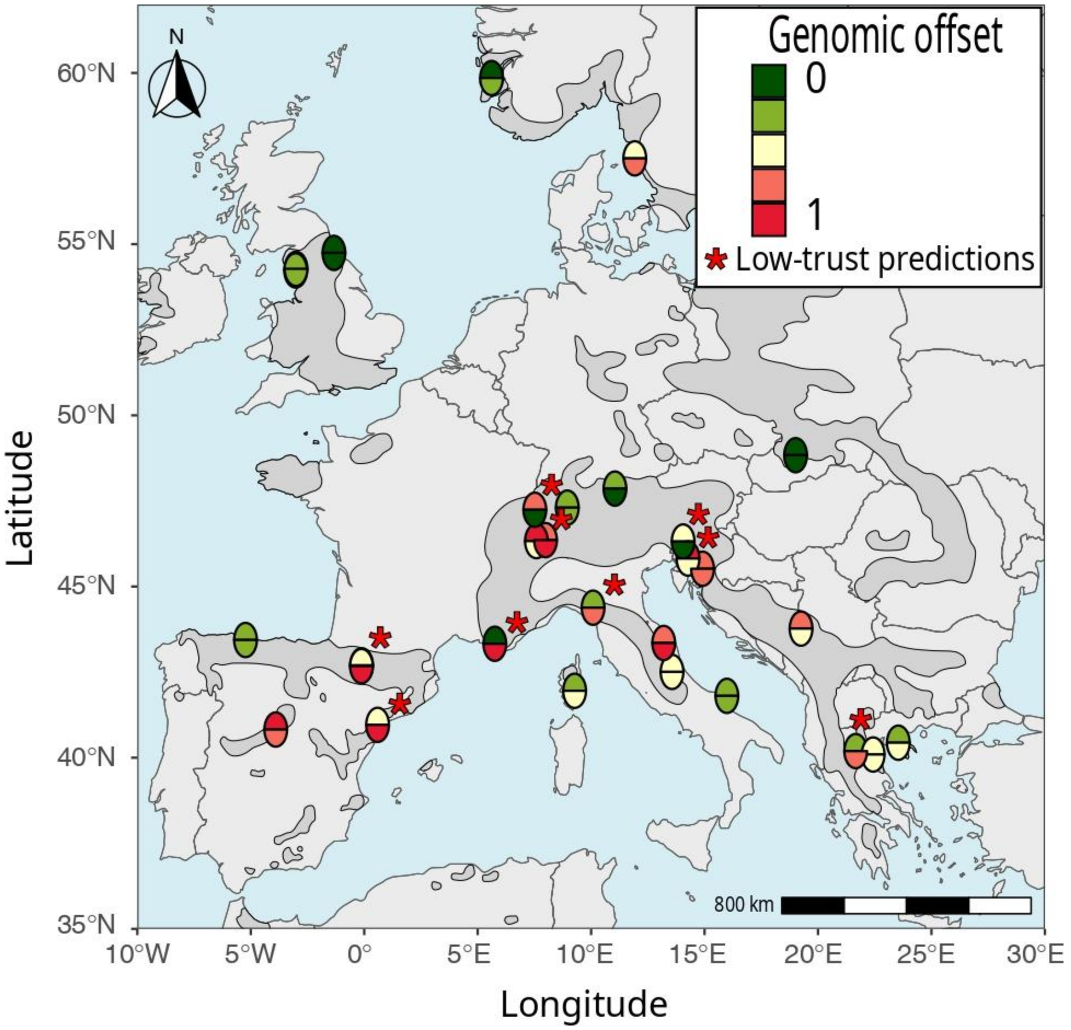


*Figure S21*. Genomic offset predictions for the RDA (lower half-circle) and GF (upper half-circle) models using the *outlier* set of SNPs for the mean climate across five GCMs for the 2041-2070 time interval under the SSP 3-7.0 using the 1901-1950 climate as reference period. Genomic offset predictions were standardised to range from zero (low - green) to one (high - red) and categorised into five classes. Red stars represent the populations where the predictions of the two models differ by more than two classes.

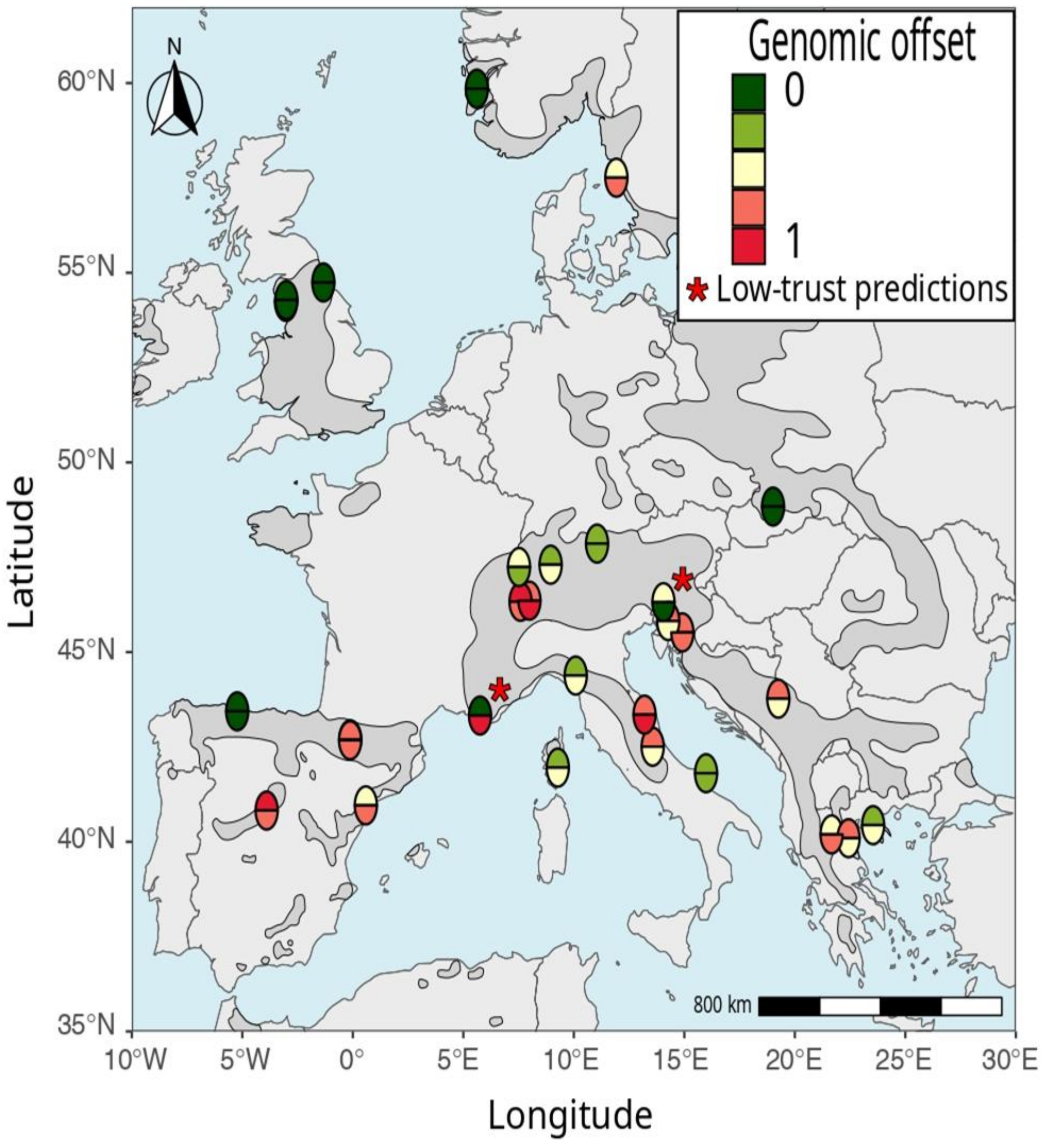


*Figure S22*. Genomic offset predictions for the RDA (lower half-circle) and GF (upper half-circle) models using the *all_outlier* set of SNPs for the mean climate across five GCMs for the 2041-2070 time interval under the SSP 3-7.0 using the 1901-1950 climate as reference period. Genomic offset predictions were standardised to range from zero (low - green) to one (high - red) and categorised into five classes. Red stars represent the populations where the predictions of the two models differ by more than two classes.
